# Morphine analgesia and μ opioid receptor signaling require programed death protein 1

**DOI:** 10.1101/580506

**Authors:** Zilong Wang, Changyu Jiang, Qianru He, Megumi Matsuda, Qingjian Han, Kaiyuan Wang, Sangsu Bang, Ru-Rong Ji

**Affiliations:** Center for Translational Pain Medicine, Department of Anesthesiology, Duke University Medical Center, Durham, North Carolina, 27710; Department of Neurobiology, Duke University Medical Center, Durham, North Carolina, 27710; Department of Cell Biology, Duke University Medical Center, Durham, North Carolina, 27710

## Abstract

Opioids such as morphine produce analgesia via mu opioid receptor (MOR), but opioid receptor signaling is not fully understood. Here we report that morphine analgesia and MOR signaling require neuronal Programmed cell death protein-1 (PD-1). We found that morphine-induced antinociception following systemic or intrathecal injection was compromised in *Pd1*^-/-^ mice. Morphine analgesia was also abrogated in wild-type mice after treatment with Nivolumab, a clinically used anti-PD-1 monoclonal antibody. Morphine produced analgesia by suppressing calcium currents in DRG neurons and excitatory synaptic transmission in spinal cord neurons, but strikingly, both actions were impaired by PD-1 blockade. In a mouse model of bone cancer, the antinociceptive action of systemic morphine was compromised in *Pd1*^-/-^ mice. Finally, PD-L1 and morphine produce synergistic analgesia. Our findings demonstrate that PD-1 also acts as a neuro checkpoint inhibitor and mediates opioid-induced analgesia and MOR signaling in neurons.

---

Programmed cell death ligand-1 (PD-L1) is an immune checkpoint inhibitor and suppresses immunity through PD-1 receptor expressed on immune cells^1-3^. Emerging immune therapies using anti-PD1 and anti-PD-L1 monoclonal antibodies have shown success in treating various cancers including melanoma by immune activation^4-6^. However, PD-1 signaling in neurons are largely unknown. We recently reported that primary sensory neurons of dorsal root ganglion (DRG) express PD-1 receptor, and activation of PD- 1 by PD-L1 inhibits neuronal excitability and pain^7^. Notably, PD-L1 is also produced by non-malignant tissues including DRG and spinal cord^7^, implicating a physiological role of PD-L1. Opioids such as morphine produce analgesia via mu opioid receptor (MOR)^8^, which is expressed in the central nervous system (brain and spinal cord) and peripheral nervous system (DRG and nerves)^9-12^. MOR mediates both analgesic and reward effects of opioids, such as morphine^8,13^. This study was undertaken to investigate the interactions between PD-1 and MOR, two inhibitory and analgesic receptors in the peripheral and central nervous system.

We first examined whether morphine analgesia would be altered in *Pd1* knockout (KO, *Pd1*^-/-^) mice after systemic injection (1, 3, and 10 mg/kg, s.c) using both tail-flick test and hot plate test. Tail-flick test in wild-type (WT) mice revealed a rapid (<0.5 h) and dose-dependent increase in tail-flick latency (TFL) for > 3 h after morphine treatment (Fig. 1A; and fig. S1A). Notably, this antinociception was compromised in KO mice, showing a shorter duration of < 3 h. Area-under-the-curve (AUC) analysis showed a 40% reduction in morphine analgesia at the dose of 10 mg/kg in KO mice, as compared to WT mice (Fig. 1A; and fig. S1A-C). Hot plate test also showed an impairment of morphine analgesia in KO mice, with a 61% reduction in AUC (Fig. 1B; and fig. S1D-F).

**Figure 1.**
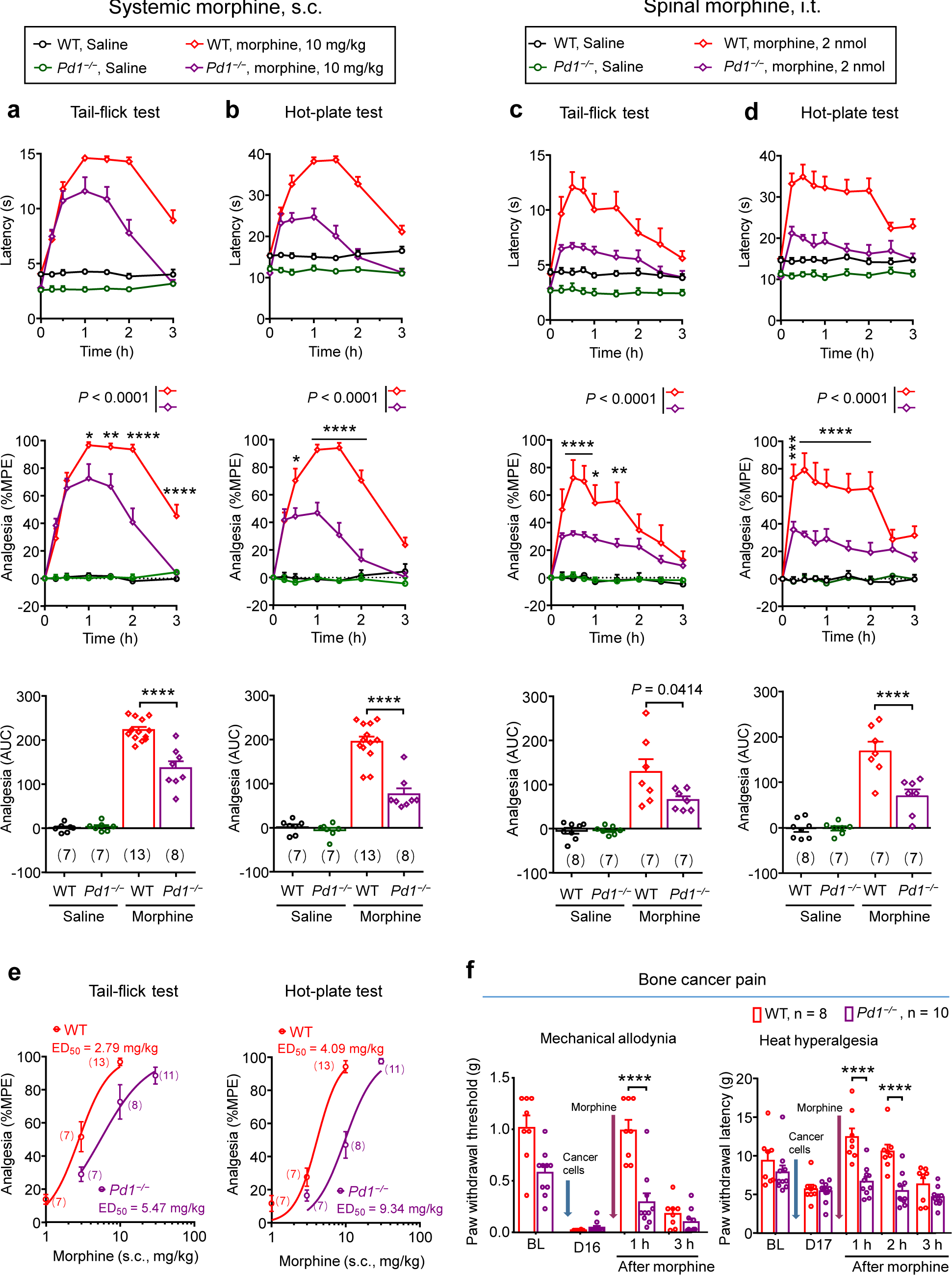
Morphine analgesia in naive and bone cancer-bearing mice is decreased after *Pd1* deletion. **(a, b)** Systemic antinociception of morphine following subcutaneous injection (s.c., 10 mg/kg) is decreased in *Pd1*^*-/-*^ mice in tail-flick (a) and hot-plate (b) tests. Upper panels, latency in both tests. Middle panels, percentage of maximum possible effect (%MPE) in both testes. \**P* < 0.05, \*\**P* < 0.01, \*\*\*\**P* < 0.0001, morphine WT vs. morphine KO, two-way ANOVA, followed by Bonferroni’s post hoc test. Bottom panels, Area Under the Curve (AUC) of %MPE, \*\*\*\**P* < 0.0001, one-way ANOVA, followed by Bonferroni’s post hoc test. *n* = 7, 7, 13, 8 mice per group, as indicated brackets. **(c, d)** Spinal antinociception of morphine following intrathecal injection (i.t., 2 nmol) is decreased in *Pd1*^*-/-*^ mice in tail-flick (c) and hot-plate (d) tests. Upper panels, latency in both tests. Middle panels, %MPE in both testes. \**P* < 0.05, \*\**P* < 0.01, \*\*\**P* < 0.001, \*\*\*\**P* < 0.0001, morphine WT vs. morphine KO, two-way ANOVA, followed by Bonferroni’s post hoc test. Bottom panels, AUC of %MPE, *P* = 0.0414 (tail-flick test), \*\*\*\**P* < 0.0001 (hot plate test), one-way ANOVA, followed by Bonferroni’s post hoc test. *n* = 8, 7, 7, 7 mice per group, as indicated brackets. **(e)** Dose-response curve of systemic antinociception of morphine (s.c., 10 mg/kg) in tail-flick (left) and hot-plate (right) tests. Note increased ED_50_ values in KO mice compared with WT mice. *n* = 7, 7, 13 in WT mice, n = 7, 8, 11 in mice, as indicated in brackets. **(f)** Spinal antinociception of morphine (i.t., 2 nmol) in bone cancer pain is decreased in *Pd1*^*-/-*^ mice. Left, mechanical allodynia, assessed in the von Frey test. Right, heat hyperalgesia assessed in the Hargreaves test. After the baseline testing, Louis lung cancer cells were inoculated into tibia (2 × 10^5^ cells in 2 µL) on Day 0 (blue arrows), and morphine was administered on Day 16 or 17 (D16 or D17, purple arrows). \*\*\*\**P* < 0.0001, two-way ANOVA, followed by Bonferroni’s post hoc test. n = 8 (WT) and 10 (KO) mice per group. Data are Mean ± SEM.

Next, we assessed the central mechanism of morphine analgesia in WT and *Pd1*^-/-^ mice. Spinal injection of morphine via intrathecal route (2 nmol, i.t.) elicited marked antinociception in tail-flick and hot plate tests in WT mice; but this spinal analgesic action of morphine was also compromised in *Pd1*^-/-^ mice (Fig. 1, C and D). Compared with WT mice, KO mice exhibited a right-shift of the dose-response curve in response to systemic morphine (Fig. 1E). Collectively, these results suggest that PD-1 is required for inducing morphine analgesia via both systemic (peripheral and central) and central (spinal) actions.

Opioids induce analgesia through different types of opioid receptors (OR), μOR (MOR), δOR (DOR), and κOR (KOR), which are expressed by DRG neurons and their central terminals as well as interneurons in the spinal cord dorsa horn^9, 13, 14^. Intrathecal injection of respective agonists of MOR (DAMGO, 0.1 nmol, i.t), DOR (DEDPE, 5 nmol, i.t.), and KOR (U69593, 20 nmol, i.t.) evoked marked analgesia in WT mice (fig. S2A-C). *Pd1*^-/-^ mice only exhibited reduced analgesia after DAMGO but not DPDPE or U69593 treatment (Fig. 2A-C), suggesting that PD-1 primarily affects the MOR-mediated analgesia.

**Figure 2.**
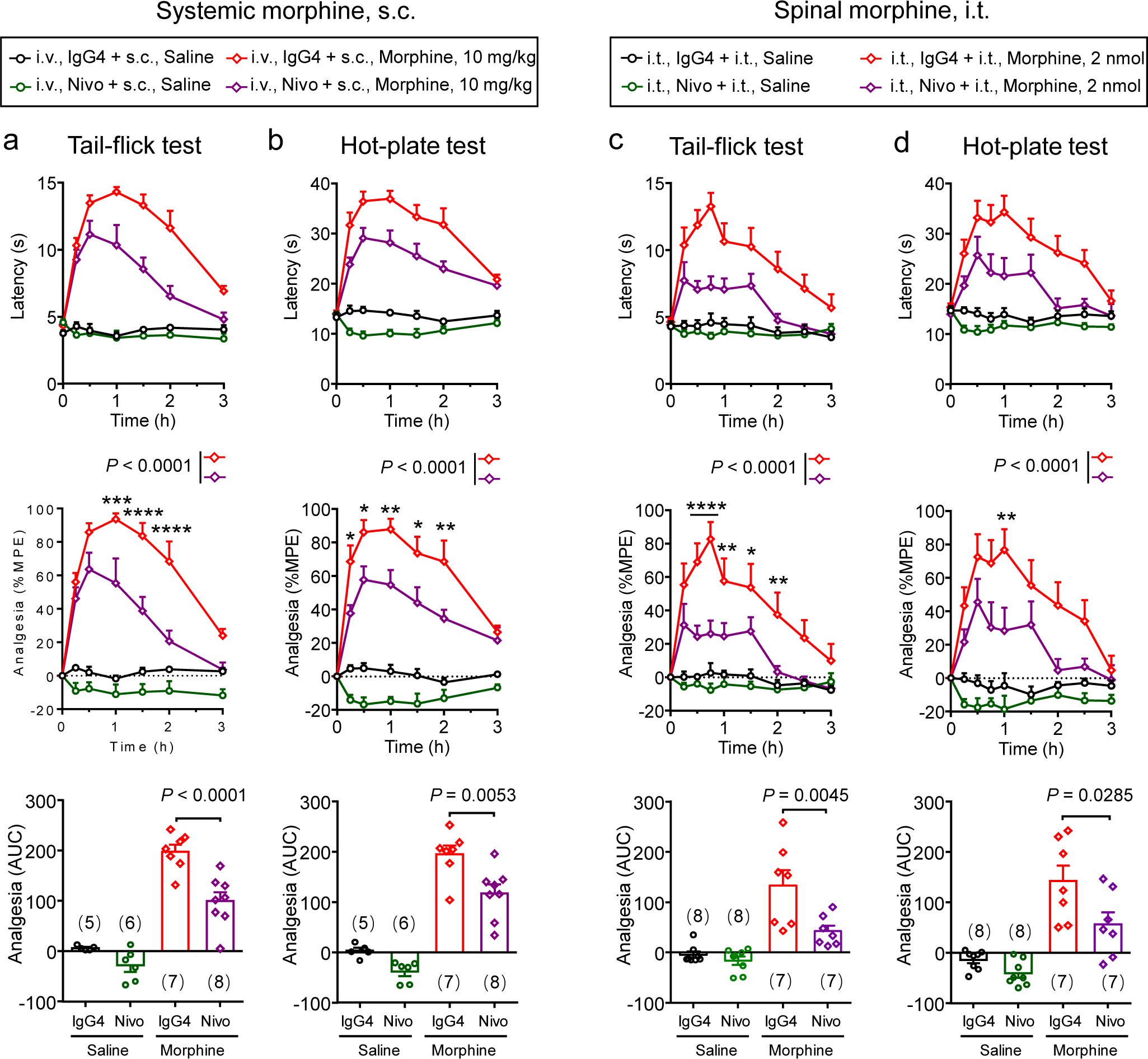
Morphine analgesia is suppressed by anti-PD-1 antibody Nivolumab. **(a, b)** Systemic antinociception of morphine following subcutaneous injection (s.c., 10 mg/kg) is decreased after intravenous pretreatment with Nivolumab (i.v., 10 mg/kg) in tail-flick (a) and hot-plate (b) tests. Upper panels, latency in both tests. Middle panels, %MPE in both testes. \**P* < 0.05, \*\**P* < 0.01, \*\*\**P* < 0.001, \*\*\*\**P* < 0.0001, morphine IgG4 vs. morphine Nivolumab (Nivo), two-way ANOVA, followed by Bonferroni’s post hoc test. Bottom panels, AUC of %MPE, *P* < 0.0001 (tail-flick test), *P* = 0.0053 (hot plate test), one-way ANOVA, followed by Bonferroni’s post hoc test. Nivolumab or control antibody (human IgG4) was i.v. injected 60 min prior to s.c. morphine injection. *n* = 5, 6, 7, 8 mice per group, as indicated in brackets. **(c, d)** Spinal antinociception of morphine following intrathecal injection (i.t., 2 nmol) is decreased after intrathecal pretreatment with Nivolumab (i.t., 1 µg) in tail-flick (c) and hot-plate (d) tests. Upper panels, latency in both tests. Middle panels, %MPE in both testes. \**P* < 0.05, \*\**P* < 0.01, \*\*\*\**P* < 0.0001, morphine IgG4 vs. morphine Nivolumab (Nivo), two-way ANOVA, followed by Bonferroni’s post hoc test. Bottom panels, AUC of %MPE; *P* < 0.0045 (tail-flick test), *P* = 0.0285 (hot plate test), one-way ANOVA, followed by Bonferroni’s post hoc test. *n* = 8, 8, 7, 7 mice per group, as indicated in brackets. Nivolumab or control antibody (human IgG4) was i.t. injected 30 min prior to i.t. morphine injection. Data are Mean ± SEM.

Opioid is a mainstay treatment for cancer pain, which often becomes unbearable after tumor metastasis to bone tissue^15^. We assessed whether morphine would attenuate cancer pain in WT and KO mice in bone cancer pain^16^. Inoculation of Lewis lung cancer cells into tibia resulted in severe cancer pain, as indicated by reductions in mechanical and thermal pain thresholds. Systemic morphine completely reversed tumor-induced mechanical allodynia and heat hyperalgesia in WT mice (Fig. 1F). Remarkably, the anti-allodynic and anti-hyperalgesic effects of morphine were largely abolished in KO mice (Fig. 1F).

Next, we tested whether loss of morphine analgesia in *Pd1*^-/-^ mice could be recapitulated by treatment of Nivolumab, a human anti-PD-1 monoclonal antibody that is also able to evoke allodynia in naive mice^7^. Tail-flick and hot plate tests showed that the antinociception induced by systemic morphine (10 mg/kg, s.c.) was abrogated by systemic pre-treatment of Nivolumab (10 mg/kg, i.v., given 1 h prior to morphine, Fig. 2, A and B). Furthermore, spinal pretreatment with Nivolumab (1 µg, i.t., given 30 min prior to morphine) also decreased morphine (2 nmol, i.t.) antinociception at spinal level (Fig. 2, C and D). Dose-response analysis revealed marked changes in ED_50_ values of morphine analgesia in KO mice and Nivolumab-treated mice, as compared to WT mice. WT mice displayed an ED_50_ value of 2.8 and 4.1 mg/kg in tail-flick test and hot plate test, respectively. The ED_50_ value increased to 5.5 (tail-flick) and 9.3 (hot-plate) in KO mice and 8.2 (tail-flick) and 8.2 (hot-plate) in Nivolumab-treated mice, respectively (fig. S3A-E). In addition to male mice, morphine antinociception was also decreased in female KO mice and blunted by Nivolumab in female WT mice (fig. S4A-D). Taken together, our data demonstrate that *Pd1* deletion and PD-1 blockade result in a substantial reduction in morphine analgesia and indicate a functional interaction between PD-1 and MOR receptors.

To determine the mechanisms by which PD-1 regulates opioid analgesia, we examined the PD-1/MOR interactions at cellular level. In DRG, MOR is mainly expressed by small-diameter nociceptive neurons as well as some medium-diameter neurons^9^, ^12^, ^17^. We assessed colocalization of *Pd1* and *Oprm1* mRNA expression in mouse DRG using a sensitive RNAscope assay. We observed high level of co-localization of *Pd1* and *Oprm1* in DRG neurons (Fig. 3, A and B). To determine the mRNA expression levels of *Pd1* and *Oprm1* in individual DRG neurons, we quantified the number of fluorescence-labeled puncta in positive neurons. We found that around 45% neurons express *Pd1* mRNA, 40% neurons express *Oprm1* mRNA, and 30% DRG neurons expressed both mRNAs (Fig. 3C). Among the *Pd1 mRNA*+ cells, around 60% of them co-expressed *Oprm1* (Fig. 3C).

**Figure 3.**
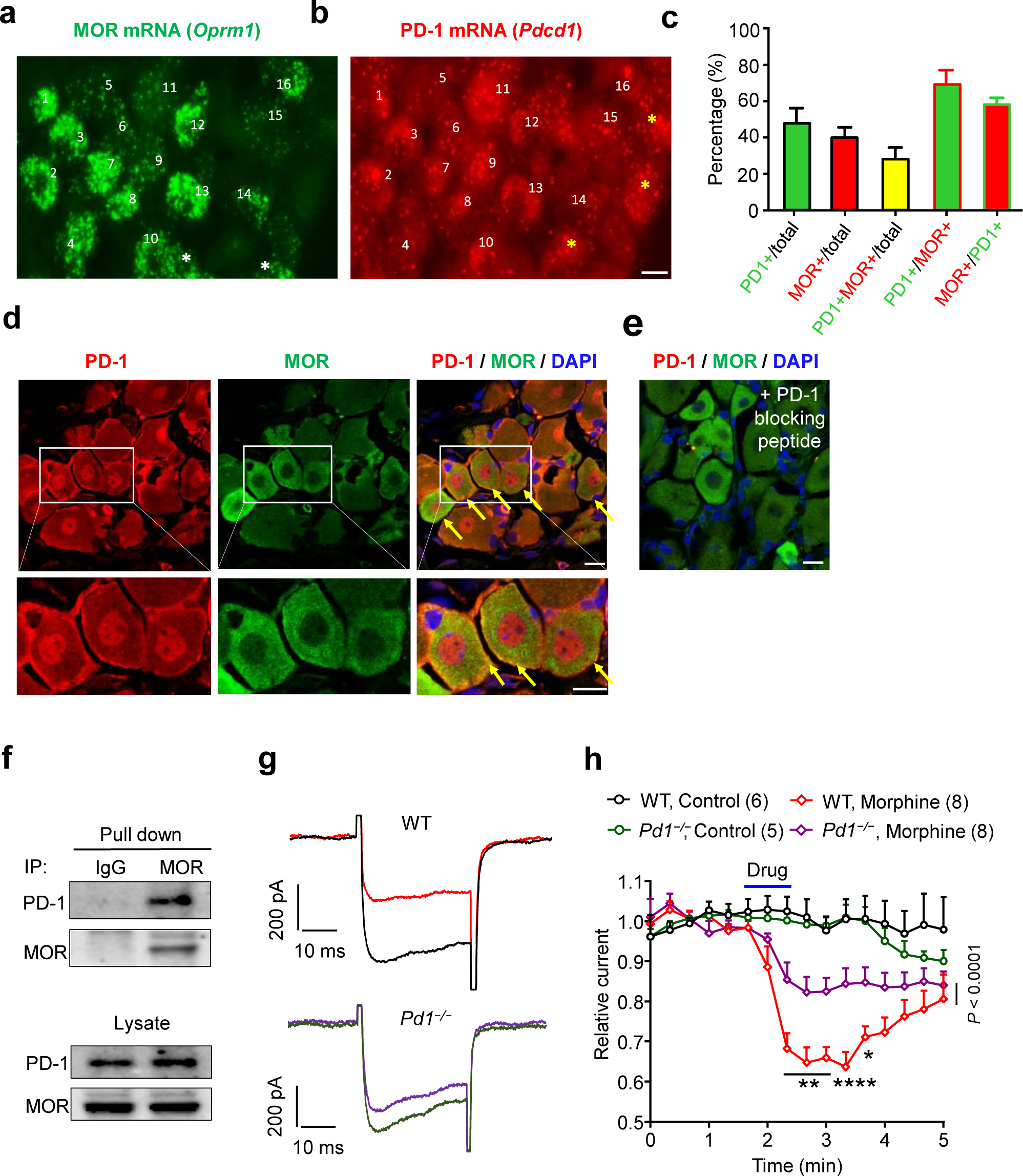
PD-1 and MOR are co-expressed in mouse DRG neurons and have functional interaction. **(a**,**b)** In situ hybridization (ISH, RNAscope) images show colocalization of *Oprm1* mRNA (a) and *Pd1* mRNA (b) in mouse DRG neurons of WT mice. Sixteen pairs of numbers show 16 neurons co-expressing both mRNAs. White starts (a) and yellow stars (b) show single-labeled neurons. Scale bar, 20 μm. **(c)** The percentage of positive neurons for *Pd1* and *Oprm* mRNA expression and co-expression in DRG. Positive cells were defined as ≥ 20 puncta in 1000 μm^2^. 1244 neurons from 3 mice were analyzed. **(d)** Double staining showing co-localization of PD-1 and MOR in DRG. Yellow arrows indicate double-labeled neurons. Low panels, enlarged boxes show co-expression of both receptors on the cell surface revealed by yellow arrows. Scale bars, 10 μm. **(e)** Absence of PD-1 immunostaining after the treatment with a PD-1 blocking peptide. Blue DAPI staining shows cell nuclei. Scale, 10 μm. **(f)** Co-IP showing PD-1/MOR interaction in DRG. DRG lysate was immunoprecipitated with control IgG or MOR antibody and immunoblotted with PD-1 or MOR antibody. This experiment was repeated 3 times. **(g, h)** Morphine-induced inhibition of calcium current is abrogated *Pd1*-deficient mouse DRG neurons. **(g)** Trace of calcium currents before and after morphine (10 μM) treatment in small-diameter DRG neurons (<25 μm) from WT and *Pd1*^*-/-*^ mice. **(h)** Time-course of relative calcium currents before and after morphine perfusion (10 μM, n = 5, 6, 8, 8 neurons per group, as indicated in the brackets). \**P* < 0.05, \*\**P* < 0.01, \*\*\*\**P* < 0.0001, KO vs. WT group, two-way ANOVA, followed by Bonferroni’s post hoc test. Data are Mean ± SEM.

Immunohistochemistry also showed high degree of co-localization of PD-1 and MOR immunoreactivity (IR) in DRG neurons (Fig. 3D). Especially, the co-localization was observed on the cell surface of small-diameter DRG neurons (Fig. 3D). The specificity of PD-1-IR was confirmed by the lack of staining after the blocking peptide treatment (Fig. 3D) and in KO mice^7^. Furthermore, we observed co-localization of PD-1 and MOR in axons of the mouse spinal nerves (fig. S5A), indicating axonal transport of both PD-1 and MOR from cell bodies to peripheral nerve axons.

We further examined possible PD-1/MOR interaction using proximity ligation assay (PLA) and co-immunoprecipitation (Co-IP). PLA analysis in DRG cultures revealed positive fluorescence signals on cell bodies and axons of DRG neurons (Fig. 3E; fig. S6), and the signal were absent after PD-1 blocking peptide treatment or by mission of primary antibody (fig. S6). Co-IP experiment demonstrated that PD-1 could be pulled down by MOR antibody from mouse DRG lysates (Fig. 3F). These results indicate a close proximity of PD-1 and MOR molecules and co-exist in a molecular complex in native sensory neurons.

Since anti-PD-1 immunotherapy has been extensively tested in humans, we also examined possible PD-1 and MOR interaction in human DRG and nerve tissues. We previously showed the presence of PD-1-IR on the surface of human DRG neurons and nerve axons of the human spinal nerve^7^. Further, we observed co-expression of PD-1 and MOR on the nerve axons of the human spinal nerve (fig. S5B). Co-IP experiment demonstrated that MOR or PD-1 antibody was able to pull down PD-1 or MOR from human spinal nerve lysates, respectively (fig. S5c). Collectively, it is suggested that PD-1 may also interact with MOR in human nerve tissue.

One mechanism for opioid to inhibit pain transmission is to suppress calcium channels in primary afferent neurons^18^. We recorded calcium currents in dissociated small-diameter DRG neurons (<25 μm) using whole-cell patch clamp ^18^. Morphine (10 μM) produced a 35% reduction in calcium currents in WT DRG neurons, but this inhibition was blunted in *Pd1*-deficient DRG neurons (Fig. 3, G and H). This result suggests that PD-1 regulates morphine analgesia in part through calcium channels.

The second mechanism for opioid analgesia is to inhibit neurotransmitter release and nociceptive synaptic transmission via presynaptic regulation in the spinal cord dorsal horn (SDH)^19-21^. Double staining revealed co-localization of PD-1 and MOR IR in SDH axons and axonal terminals (fig. S7A). PD-1 expression by SDH presynaptic terminals was also confirmed by its colocalization with CGRP, a neuropeptide derived from primary afferents (Fig. S7B). We prepared spinal cord slices to record spontaneous excitatory post-synaptic currents (sEPSCs) in out lamina II (IIo) neurons, which are predominantly excitatory and form a nociceptive circuit with primary C-afferents and lamina I project neurons^22, 23^. Bath perfusion of morphine (10 μM) in spinal cord slices of WT mice resulted in a marked inhibition of sEPSC frequency (44.8% inhibition, Fig. 4, A-E). However, in *Pd1*-deficient SDH neurons, this inhibition was significantly lower (22.7%, *P* = 0.0068, two-tailed t-test, KO vs. WT mice, Fig. 4E). Morphine (10 μM) also caused a mild inhibition of sEPSC amplitude (10.7%) in WT mice (*P* = 0.0068, two-tailed t-test) and had no inhibition of the sEPSC amplitude in KO mice (*P* = 0.1272, paired two-tailed t-test). No significant difference was observed in the sEPSC amplitude between WT and KO mice after morphine treatment (n=13-14 neurons, *P*= 0.2692, paired two-tailed t-test, Fig. 4E). A dose-response analysis showed that morphine suppressed sEPSC frequency at IC_50_ = 1.07 μM in WT mice, but the IC_50_ value was increased to 15.24 μM in KO mice (fig. S8A-C). A specific involvement of MOR was confirmed by a reduction of DAMGO’s (0.5 μM) inhibition of sEPSCs in KO mice (fig. S8D,E). These results suggest that *Pd1* deletion reduces opioid’s efficacy in suppressing spinal nociceptive synaptic transmission.

**Figure 4.**
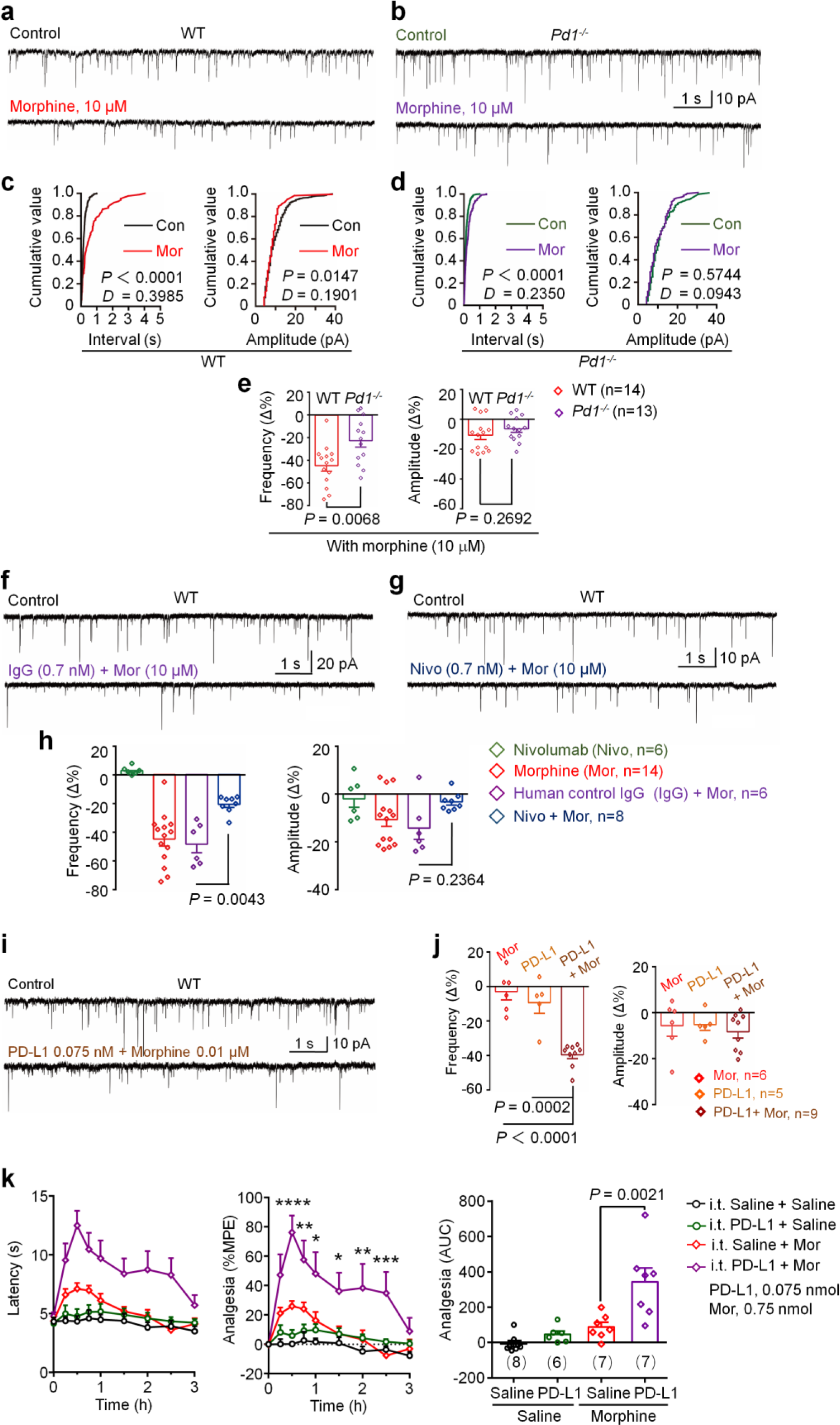
Morphine’s on spinal excitatory synaptic transmission in WT and *Pd1*^*-/-*^mice and synergistic antinociception of PD-L1 and morphine in WT mice. **(a-e)** Morphine’s suppression of spinal excitatory synaptic transmission is compromised after *Pd1* deficiency. (**a, b**) Recording traces of spontaneous excitatory postsynaptic currents (sEPSCs), without morphine (upper) or with morphine (lower, 10 μM), in lamina II neurons of spinal cord slices from WT (a) and *Pd1*^*-/-*^mice (b). (**c, d**) Cumulative histograms of the inter-event interval and amplitude of sEPSCs in WT (c) and KO (d) mice. The histograms were examined for 1 min before morphine treatment (338 events in WT mice and 415 events in KO mice) and 1 min after morphine washout (85 events in WT mice and 235 events in KO mice). **(c)** In WT neurons morphine shifted the inter-event interval and amplitude to a longer (*P* < 0.0001; Kolmogorov-Smirnov test, left) and smaller ones (*P* = 0.0147; Kolmogorov-Smirnov test, right). Traces in a and c were obtained from the same neuron. **(d)** In *Pd1* deficient neurons, the inter-event interval shift is also significant (*P* < 0.0001; Kolmogorov-Smirnov test) after morphine treatment. Traces in b and d were obtained from the same neuron. *D* indicates distance between two curves. (**e**) Percentage changes in sEPSCs frequency (left) and amplitude (right) in WT and KO mice. Morphine-induced reduction in sEPSCs frequency was impaired in *Pd1*^*-/-*^mice, compared with WT mice (*P*= 0.0068, unpaired two-tailed t-test, n=13-14 per group). There was no significant difference in sEPSCs amplitude between WT and *Pd1*^*-/-*^mice (n=13-14 per group, *P* = 0.2692, unpaired two-tailed t-test). **(f-h)** Nivolumab blocks the morphine’s inhibition of sEPSCs in spinal lamina II neurons of WT mice. (**f, g**) Recording traces of sEPSCs in spinal slices treated with morphine and control IgG (f) or treated with morphine and Nivolumab (g). (**h**) sEPSCs frequency (left) and amplitude (right) after perfusion with Nivolumab (0.7 nM) or control IgG4 together with morphine (10 μM). *P* = 0.0043, one-way ANOVA, followed by Bonferroni’s post hoc test, n = 6-14 cells per group. **(i, j)** Synergistic action of PD-L1 and morphine in suppressing sEPSCs in lamina II neurons of WT mice. (**i**) Recording traces of sEPSCs before (upper) and after (low) treatment of morphine plus PD-L1 (0.075 nM). The upper and lower traces were from same neuron. (**j**) sEPSCs frequency (left) and amplitude (right). *P*< 0.0001 or *P*= 0.0002, one-way ANOVA, n=5-9 cells per group. (**k**) Tail-flick test showing that a combination of PD-L1 and morphine injection produces synergistic antinociception. Left, withdrawal latency following i.t. injection of morphine (0.75 nmol), PD-L1 (0.01 nmol), and PD-L1 plus morphine. PD-L1 or saline was i.t. injected 30 min prior to morphine injection. Middle, %MPE, \**P* < 0.05, \*\**P* < 0.01, \*\*\**P* < 0.0001, \*\*\*\**P* < 0.0001, vs. saline group, two-way ANOVA. Right, AUC of %MPE. *P* = 0.0021, n = 8, 6, 7, 7 mice per group (indicated in brackets). Data are Mean ± SEM.

PD-L1 is a major ligand of PD-1 and produces antinociception in mouse models of pathological pain^7^. We investigated whether PD-L1 would modulate synaptic transmission via PD-1 in SDH neurons. Dose response analysis showed that the IC_50_ for PD-L1 and morphine to inhibit sEPSC frequency is 0.60 nM and 1.7 µM, respectively (fig. S9A,B), suggesting that PD-L1 is 1700 times more potent than morphine. As expected, the effects of PD-L1 on sEPSCs were blunted in *Pd1*-deficient neurons (fig. S9C). Notably, in naïve animals, *Pd1* deficiency lead to an increase in sEPSC frequency but not amplitude (fig. S9D), indicating that PD-1 is an endogenous negative regulator of synaptic transmission.

The third mechanism of opioid analgesia is to regulate potassium channels to generate outward currents in neurons^24^. Morphine (10 μM) evoked outward currents in 36.7% (5/14) SDH neurons of WT mice, and the average amplitude of outward currents was 15.8 ± 1.5 pA. In contrast, morphine-evoked outward currents were significantly smaller in KO mice (7.8 ± 1.8 pA; *P* = 0.0104, unpaired two-tailed t-test, vs. WT), despite similar response rate of 30.7% (4/13) (fig. S10A-C). PD-L1 also evoked dose-dependent outward currents (5-18 pA) in 20-50% SDH neurons, and these outward currents were abolished in KO mice (fig. S10D,E).

As expected, Nivolumab also blocked the morphine’s effects on SDH neurons in WT mice. Perfusion of spinal cord slices with Nivolumab at a very low concentration (100 ng/ml, i.e. 0.7 nM) significantly reduced the morphine’s inhibition of sEPSC frequency (*P* < 0.001, compared with human IgG4 control, Fig. 4, F-H). Nivolumab at this low concentration (0.7 nM) did not affect sEPSC frequency, but increased sEPSC frequency at high concentrations (2.1 and 10.5 nM, fig. S11A-F). Neither did the control IgG4 affect the morphine’s inhibition of sEPSC (Fig. 4H). Nivolumab also suppressed morphine’s inhibition of sEPSCs amplitude (Fig. 4H).

Finally, we assessed whether PD-L1 and morphine would produce additive or synergistic actions in inhibiting synaptic transmission and pain. At very low concentrations, PD-L1 (0.075 nM) and morphine (0.01 μM) each produced mild inhibition of sEPSC frequency (5% and 8%, respectively), but co-application of PD-L1 and morphine at these low concentrations produced a much greater inhibition (40%) of sEPSCs, suggesting a synergistic effect of these two compounds at low concentrations (Fig. 4, I and J). This synergistic action of morphine and PD-L1 was also evident in behavioral testing. Intrathecal PD-L1 (0.075 nmol) alone failed to produce any analgesia, and intrathecal morphine (0.75 nmol) only produce very mild and transient analgesia, but intrathecal co-application of PD-L1 and morphine at these low doses produced a much greater analgesia, indicating a synergistic analgesic effect (Fig. 4K and fig. S12).

In summary, we have demonstrated that the morphine’s key analgesic actions, such as antinociception in tail-flick and hot plate tests, anti-hyperalgesic and anti-allodynic effects in bone cancer pain, suppression of calcium currents in DRG neurons, as well as inhibition of sEPSCs and induction of outward currents in SDH neurons are all abrogated in *Pd1*^-/-^ mice. Co-IP suggested that PD-1 may interact with MOR in mouse DRG neurons and human nerve axons, although future studies are required to determine the details of the interaction. The functional interactions only occur between PD-1 with MOR, but not with DOR or KOR. Importantly, the defects in morphine signaling in *Pd1*^-/-^ mice can be recapitulated by Nivolumab treatment in adult mice. In *Pd1*-deficinet mice, morphine’s analgesia in bone cancer pain is largely impaired. Our findings have clinical implication, because 1) Nivolumab has been approved for treating various cancers and 2) opioids are mainstay treatments for cancer pain and peripheral analgesic actions of opioids are well appreciated^10, 12, 25, 26^. Thus, anti-PD-1 immune therapy may interfere opioid analgesia in cancer patients via disrupting the PD-1-MOR interaction in the peripheral nerve and DRG tissue. Opioid receptor has also been implicated in placebo analgesia27, and it will be interesting to know if immune therapy would interrupt this unique type of analgesia. On the other hand, PD-L1 might be used to treat clinical pain and enhance opioid analgesia in cancer and non-cancer patients.

## Supporting information

Wang_supplemental material

## Acknowledgements

This study is supported by NIH R01 grants DE17794 and DE22743 to R.-R. J.

## Author contributions

Z.W. conducted behavioral tests and electrophysiology in DRG neurons; C.J. conducted electrophysiology in spinal cord slices; Q-R.H conducted immunohistochemistry; M.M. conducted *in situ* hybridization; Q.H. and S.B. conducted immunoprecipitation; K.W. prepared mouse model of bone cancer. R.-R. J. supervised the project; R.-R. J., Z.W., and C. J. wrote the paper.

## References

1. Nishimura, H., Nose, M., Hiai, H., Minato, N. & Honjo, T. Development of lupus-like autoimmune diseases by disruption of the PD-1 gene encoding an ITIM motif-carrying immunoreceptor. Immunity 11, 141–151 (1999).

2. Freeman, G.J., et al. Engagement of the PD-1 immunoinhibitory receptor by a novel B7 family member leads to negative regulation of lymphocyte activation. The Journal of experimental medicine 192, 1027–1034 (2000).

3. Gordon, S.R., et al. PD-1 expression by tumour-associated macrophages inhibits phagocytosis and tumour immunity. Nature 545, 495–499 (2017).

4. Herbst, R.S., et al. Predictive correlates of response to the anti-PD-L1 antibody MPDL3280A in cancer patients. Nature 515, 563–567 (2014).

5. Brahmer, J.R., et al. Safety and activity of anti-PD-L1 antibody in patients with advanced cancer. N.Engl.J Med. 366, 2455–2465 (2012).

6. Topalian, S.L., et al. Safety, activity, and immune correlates of anti-PD-1 antibody in cancer. N.Engl.J Med. 366, 2443–2454 (2012).

7. Chen, G., et al. PD-L1 inhibits acute and chronic pain by suppressing nociceptive neuron activity via PD-1. Nature neuroscience (2017).

8. Matthes, H.W., et al. Loss of morphine-induced analgesia, reward effect and withdrawal symptoms in mice lacking the mu-opioid-receptor gene. Nature 383, 819–823 (1996).

9. Ji, R.R., et al. Expression of mu-, delta-, and kappa-opioid receptor-like immunoreactivities in rat dorsal root ganglia after carrageenan-induced inflammation. J.Neurosci. 15, 8156–8166 (1995).

10. Stein, C., et al. Peripheral mechanisms of pain and analgesia. Brain Res.Rev. 60, 90–113 (2009).

11. Mousa, S.A., Zhang, Q., Sitte, N., Ji, R. & Stein, C. beta-Endorphin-containing memory-cells and mu-opioid receptors undergo transport to peripheral inflamed tissue. J.Neuroimmunol. 115, 71–78 (2001).

12. Scherrer, G., et al. Dissociation of the opioid receptor mechanisms that control mechanical and heat pain. Cell 137, 1148–1159 (2009).

13. Corder, G., Castro, D.C., Bruchas, M.R. & Scherrer, G. Endogenous and Exogenous Opioids in Pain. Annu Rev Neurosci (2018).

14. Chavkin, C., James, I.F. & Goldstein, A. Dynorphin is a specific endogenous ligand of the kappa opioid receptor. Science 215, 413–415 (1982).

15. Mantyh, P.W. Cancer pain and its impact on diagnosis, survival and quality of life. Nat.Rev.Neurosci 7, 797–809 (2006).

16. Wakabayashi, H., et al. Decreased sensory nerve excitation and bone pain associated with mouse Lewis lung cancer in TRPV1-deficient mice. J Bone Miner Metab 36, 274–285 (2018).

17. Guan, J.S., et al. Interaction with vesicle luminal protachykinin regulates surface expression of delta-opioid receptors and opioid analgesia. Cell 122, 619–631 (2005).

18. Andrade, A., Denome, S., Jiang, Y.Q., Marangoudakis, S. & Lipscombe, D. Opioid inhibition of N-type Ca2+ channels and spinal analgesia couple to alternative splicing. Nature neuroscience 13, 1249–1256 (2010).

19. Drdla-Schutting, R., Benrath, J., Wunderbaldinger, G. & Sandkuhler, J. Erasure of a spinal memory trace of pain by a brief, high-dose opioid administration. Science 335, 235–238 (2012).

20. Chen, G., et al. beta-arrestin-2 regulates NMDA receptor function in spinal lamina II neurons and duration of persistent pain. Nat Commun 7, 12531 (2016).

21. Heinke B G.E., and Sandkuhler J. Multiple Targets of mu-Opioid Receptor-Mediated Presynaptic Inhibition at Primary Afferent A{delta}- and C-Fibers. in J.Neurosci. 2011;31 1313-1322 (2011).

22. Todd, A.J. Neuronal circuitry for pain processing in the dorsal horn. Nat.Rev.Neurosci. 11, 823–836 (2010).

23. Duan, B., Cheng, L. & Ma, Q. Spinal Circuits Transmitting Mechanical Pain and Itch. Neuroscience bulletin 34, 186–193 (2018).

24. North, R.A. & Williams, J.T. On the potassium conductance increased by opioids in rat locus coeruleus neurones. J Physiol 364, 265–280 (1985).

25. Levine, J.D. & Taiwo, Y.O. Involvement of the mu-opiate receptor in peripheral analgesia. Neuroscience 32, 571–575 (1989).

26. Spahn, V., et al. A nontoxic pain killer designed by modeling of pathological receptor conformations. Science 355, 966–969 (2017).

27. Levine, J.D., Gordon, N.C. & Fields, H.L. The mechanism of placebo analgesia. Lancet 2, 654–657 (1978).

