## Supplementary material for "Morphine analgesia and μ opioid receptor signaling require programed death protein 1": Wang_supplemental material

**This file includes:**

Materials and Methods
Figs. S1 to S12

### Materials and Methods

Reagents

Mouse PD-L1 (Catalog: ab130039) and human IgG4 (Catalog: ab90286) were purchased from Abcam. Nivolumab (OPDIVO®), a humanized anti-PD-1 antibody, was from Bristol-Myers Squibb. Morphine Sulfate was obtained from WEST-WARD pharmaceuticals. DAMGO (Catalog: E7384), DPDPE (Catalog: E3888), and U69593 (Catalog: U103) were obtained from Sigma Aldrich.

Animals

*Pd1* (Pdcd1) knockout mice with a C57BL/6 background were purchased from the Jackson Laboratory (stock no.: 021157) and maintained at the Duke animal facility. Young mice (5–7 weeks of both sexes) were used for electrophysiological studies in the spinal cord and DRG neurons. Adult male mice (8-10 weeks), including knockout mice and corresponding wild-type control mice, as well as some CD1 mice (Charles River), were used for behavioral and pharmacological studies. Female mice were also used for behavioral test in Fig. S4. Mice were group-housed on a 12-hour light/12-hour dark cycle at 22 ± 1 °C with free access to food and water. Animals were randomly assigned to each group. Sample sizes were estimated based on our previous studies for similar types of behavioral, biochemical, and electrophysiological analyses (1-3). Two to five mice were housed in each cage. Animal experiments were conducted in accordance with the National Institutes of Health Guide for the Care and Use of Laboratory Animals.

Mouse bone cancer pain model and behavioral testing

Lewis lung carcinoma cell line (LLC1) was obtained from ATCC. Before the inoculation, the cancer cells were digested with 0.25% trypsin and made into a suspension of 1×108/ml cells in PBS. The inoculation was performed as previously described (4). Briefly, mice were anesthetized with 3% isoflurane, and a 0.5-1cm superficial incision was made near the knee joint to expose the patellar ligament. Then a 25-gauge needle was inserted at the site of the intercondylar notch of the left femur into the femoral cavity. The needle was connected with a 10 µL microinjection syringe containing 2 µL suspension of $2 \times 10^5$ tumor cells, as well as 2 µL absorbable gelatin sponge solution for the closure of the injection site. The contents of the syringe were slowly injected into the femoral cavity for a duration of 2 min. To further prevent leakage of tumor cells, the outside injection site was sealed with silicone adhesive (Kwik-Sil, World Precision Instruments). The wound was then closed with silk sutures and re-swabbed.

Drug injection. For intravenous injection, anti-PD-1 antibody (Nivolumab, 10 mg/kg in 100 µl saline) or control antibody (human IgG4) was administered into the tail vein of mouse. For intrathecal injection, spinal cord puncture was made by a Hamilton microsyringe (Hamilton) with a 30-G needle between the L5 and L6 level to deliver reagents (5 µl) to the cerebral spinal fluid(3). For subcutaneous injection, morphine (1, 3, 10, 30 mg/kg in 250 µl saline) was injected on the back with a 30-G needle.

Behavioral testing.

All animals were habituated to testing environment for at least 2 days before the baseline testing. All the tests were concluded blindly.

Tail-flick test. Mice were gently held by hand with a terry glove. The exposed distal part of the tail (3 cm) was immersed into 50°C hot water. The tail-flick latency was defined as the time required for a mouse to flick or remove its tail out of hot water. A maximum cut-off value of 15 seconds was set to avoid thermal injury. Tail-flick latency was assessed before and after drug injection. Data are also expressed as the maximum possible effect (MPE), calculated as MPE (%) = 100 × [(postdrug response – baseline response)/(cutoff response − baseline response)]. The MPE (%) data from each animal were also converted to area under the curve (AUC).

Hot-plate test. For all the experiments, hot-plate test was conducted after tail-flick test. Mice were placed on the hot plate at 53 °C and the reaction time was scored when the animal began to exhibit signs of pain avoidance such as jumping or paw licking. A maximum cut-off value of 40 seconds was set to avoid thermal injury. Data are expressed as the maximum possible effect (MPE) calculated as MPE (%) = 100 × [(postdrug response – baseline response)/(cutoff response − baseline response)]. The MPE (%) data from each animal were converted to area under the curve (AUC).

Mouse bone cancer pain testing. Mechanical and thermal sensitivity was assessed by von Frey test and Hargreaves test, respectively. For Von Frey test, animals were habituated to the testing environment daily for at least 2 d before baseline testing. Animals were confined in boxes placed on an elevated metal mesh floor and the hind paws were stimulated with a series of von Frey hairs with logarithmically increasing stiffness (0.02–2.56g, Stoelting), presented perpendicularly to the central plantar surface. We determined the 50% paw withdrawal threshold by the up-down method. Thermal sensitivity was tested using Hargreaves radiant heat apparatus (IITC Life Science). The basal paw withdrawal latency was adjusted to 9–12 s, with a cutoff of 20 s to prevent tissue damage.

In situ hybridization (ISH)

Animals were deeply anesthetized with isoflurane and transcardially perfused with PBS, followed by 4% paraformaldehyde. After the perfusion, the L4-L5 DRGs were removed and postfixed in the same fixative for 2 hours at 4 ˚C. Then, the tissues were cryopreserved in 30% sucrose/ PBS solution for 2 days. DRG sections (12 µm) were cut using a cryostat. In situ hybridization was performed using RNAscope system (Advanced Cell Diagnostics) according to the manufacturer’s protocol. Pretreatment consisted of dehydration, followed by incubation with hydrogen peroxide and protease IV at room temperature. Subsequently, the protocol for the Multiplex Fluorescent Kit v2 was followed using commercial probes for PD-1 (Mm-Pdcd1, #416781) and MOR (Mm-Oprm1-C3, # 315841-C3). ISH images were captured using a Nikon fluorescence microscope. For quantification, all the images were taken using the same acquisition settings, and five DRG sections from each animal were selected and four animals were included for data analysis. To determine the percentage of labeled cells, we quantified the number of fluorescence puncta using the bioimage analysis software QuPath. Borders were drawn manually around neurons to define regions of interest. The ‘subcellular detection’ analysis feature of QuPath was used to count individual spots. The cells with more than 20 puncta per area (1000 µm2) were classified as positive neurons. Notably, the background (DRG tissue without cells) is less than 1 puncta per area. The percentage of positive neurons could be much higher when the stringency for defining positive cells is lower (e.g., 5 or 10 puncta per area).

Immunohistochemistry in mouse and human tissues

Mice were deeply anesthetized with isoflurane and perfused through the ascending aorta with PBS, followed by 4% paraformaldehyde. After the perfusion, L4-L5 DRGs, L4-L5 spinal cords, and L4-L5 spinal nerves were removed from the mice and post-fixed in the same fixative overnight. Human DRG-spinal nerve tissues (L4-L5) were from 3 disease-free donors from NDRI (National Disease Research Interchange) and the DRG attached spinal nerves were cut and immediately post-fixed in 4% paraformaldehyde overnight. The samples were then dehydrated with 30% sucrose solution, imbedded in Tissue-Tek O.C.T., and sliced into sections (14 μm for DRG and spinal nerve sections, 30 μm for free floating spinal cord sections) in a cryostat. The sections were blocked with 5% goat serum for 1 h at room temperature and then incubated overnight at 4 °C with the following primary antibodies: anti-PD-1 antibody (rabbit, 1:300, Sigma, catalog: PRS4065), anti-MOR antibody (guinea pig, 1:100, Neuromics, catalog: GP10106), anti-CGRP antibody (goat, 1:500, Bio-rad, catalog: 1720-9007). The sections were washed in PBS and incubated with the following secondary antibodies (1:400, Jackson Immunoresearch) for 2 h at room temperature: Cy3-donkey anti-rabbit (catalog: 711-165-152), FITC-donkey anti-goat (catalog: 705-095-003) and Alexa Fluor® 488 -goat anti guinea pig (catalog: 106-545-003). DAPI (1:1,000, Thermo Scientific, catalog: 62248) was used to stain the cell nuclei in tissue sections. The sections were then washed with PBS and mounted in fluorescent mounting medium, and observed under a confocal laser scanning microscope (SP5 Inverted confocal-LSRC, Leica Microsystems). To confirm the specificity of PD-1 antibody, blocking experiments were performed in DRG, spinal cord and spinal nerve sections by using a mixture of anti-PD-1 antibody (1:300) and PD-1 Blocking Peptide (1:300, Sigma, catalog: SBP4065), as we previously demonstrated (5).

Proximity ligation assay (PLA)

PLA was performed using Duolink reagents (Sigma-Aldrich, catalog: DUO92101) on cultured mouse DRG neurons. The dissociated DRG neurons were cultured as described above. Two days later after culture, cells were fixed with 4% paraformaldehyde for 15 min at room temperature and permeabilizated with 0.1% Triton X-100 for 30 min. PLA conducted according to the Duolink PLA Protocol from Sigma-Aldrich to examine possible interaction of PD-1 and MOR. Briefly, cells on coverslips were blocked with Duolink Blocking solution and incubated with a mixture of two primary antibodies (1:100 rabbit anti-PD-1, catalog: PRS4065, Sigma; 1:200 mouse anti-MOR, mAb63256, Abcam) overnight at 4°C. After incubation, the coverslips were washed with wash buffer A and then incubated with secondary antibodies (anti-mouse MINUS probe and anti-rabbit PLUS probe) for 1 h at 37 °C. Coverslips were then washed with wash buffer A and incubated with PLA ligase in ligation buffer for 30 min at 37 °C. After incubation, coverslips were washed with wash buffer A and incubated with the polymerase in amplification buffer for 100 min at 37 °C, then washed with wash buffer B followed with 0.01 × wash buffer B for 1 min. Coverslips were mounted in mounting medium containing DAPI (Sigma-Aldrich) and images were captured using a Nikon fluorescence microscope. To confirm the specificity of the PD-1 antibody, blocking experiments were performed by adding the PD-1 antibody blocking peptide (1:200, Sigma, catalog: SBP4065). The negative control was conducting by omission of primary antibody.

Immunoprecipitation

Mouse DRG tissues were lysed in ice-cold immunoprecipitation buffer (10 mM Tris-HCl, pH 7.4, 150 mM NaCl, 1 mM EDTA, 1% Triton X-100 and 10% glycerol). The lysates were immunoprecipitated with 0.5 mg of anti-MOR antibody (Neuromics, GP10106) and then incubated with protein G-agarose beads (Roche). Then the beads were collected and washed at least three times with the immunoprecipitation buffer. For immunoblotting, the lysates or beads were incubated in SDS-PAGE loading buffer for 30 min at 50°C, supernatant was collected after centrifugation at 13000 rpm. The samples were separated on an SDS-PAGE gel, transferred, and probed with anti-PD1 antibody (1:1000; Sigma, PRS4065) and anti-MOR antibody (1:1000, Abcam, ab10275). The immunoreactive bands were detected with horseradish peroxidase-conjugated secondary antibody, visualized with enhanced chemiluminescence (Thermo scientific, Pittsburgh, PA). Each experiment was repeated three times.

Whole-cell patch clamp recording in cultured DRG neurons

DRGs were removed aseptically from mice (5-8 weeks) and incubated with collagenase A (0.2 mg/ml, Roche)/dispase-II (3 mg/ml, Roche) at 37°C for 90 min, then the cells were mechanically dissociated with a flame polished Pasteur pipette in the presence of 0.05% DNAse I (Sigma). DRG cells were plated on 0.1 mg/ml poly-D-lysine–coated glass coverslips and grown in the neurobasal medium supplemented with 10% FBS, 2% B-27 supplement, and 1% penicillin/streptomycin. DRG neurons were grown for 24-48 hours before use. Whole-cell patch clamp recordings were performed in small-diameter neurons (<25 μm) at room temperature using an Axopatch-200B amplifier (Axon Instruments) with a Digidata 1440A (Axon Instruments). The patch pipettes were pulled from borosilicate capillaries (World Precision Instruments, Inc.) using a P-97 Flaming/Brown micropipette puller (Sutter Instrument Co.). Pipette resistance was 4-6 MΩ for whole-cell recording. Calcium current was evoked by a 40-ms step depolarization to -10 mV from -80 mV, as previously reported(6). The external solution contained 135 mM TEA-Cl, 1 mM CaCl2, 10 mM HEPES, 4 mM MgCl2 and 0.1 μM TTX, adjusted to a pH of 7.4 with TEA-OH. The pipette solution contained 126 mM CsCl, 5 mM Mg-ATP, 10 mM EGTA and 10 mM HEPES, adjusted to a pH of 7.3 with CsOH.

Whole-cell patch clamp recording in spinal cord slices

Mice of both sexes (5-7 weeks old) were anesthetized with urethane (1.5-2.0 g/kg, i.p.). The lumbosacral spinal cord was quickly removed and placed in ice-cold sucrose-ACSF which was saturated with 95% O2 and 5% CO2 and maintained at room temperature. After spinal extraction under anesthesia, animals were sacrificed by decapitation. Transverse spinal slices (300-400 μm) were prepared using a vibrating microslicer (VT1200s Leica). The slices were incubated at 32℃ for at least 30 min in regular ACSF (NaCl 126 mM, KCl 3 mM, MgCl2 1.3 mM, CaCl2 2.5 mM, NaHCO3 26 mM, NaH2PO4 1.25 mM and glucose 11 mM), equilibrated with 95% O2 and 5% CO2.  The slices were placed in a recording chamber and completely submerged and perfused at a flow rate of 1.5-3 ml/min with ACSF which was saturated with 95% O2 and 5% CO2 and maintained at room temperature. Lamina II neurons in lumbar segments were identified as a translucent band under a microscope (BX51WIF; Olympus) with light transmitted from below. Whole-cell voltage-clamp recordings were made from outer lamina II neurons by using patch-pipettes fabricated from thin-walled, fiber-filled capillaries. Patch-pipette solution used to record spontaneous excitatory postsynaptic currents (sEPSCs) contained: K-gluconate 135 mM, KCl 5 mM, CaCl2 0.5 mM, MgCl2 2 mM, EGTA 5 mM, HEPES 5 mM, Mg-ATP 5 mM (pH 7.3 adjusted with KOH, 300mOsm). The patch-pipettes had a resistance of 8–10 M. The sEPSCs recordings were made at a holding potential (VH) of -70 mV in the presence of 10 μM picrotoxin and 2 μM strychnine. Signals were acquired using an Axopatch 700B amplifier. The data were stored and analyzed with a personal computer using pCLAMP 10.3 software. sEPSC events were detected and analyzed using Mini Analysis Program ver. 6.0.3. Numerical data are given as the mean ± SEM. Statistical significance was determined as $P < 0.05$ using Student's t test. In all cases, n refers to the number of the neurons studied. All drugs were bath applied by gravity perfusion via a three-way stopcock without any change in the perfusion rate.

Statistical analyses

All data were expressed as mean ± s.e.m, as indicated in the figure legends. Statistical analyses were completed with Prism GraphPad 6.1. Behavioral data were analyzed using two-tailed student's t-test (two groups), One-Way or Two-Way ANOVA (repeated measures over a time course) followed by post-hoc Bonferroni test. Electrophysiological data were tested using one-way ANOVA (for multiple comparisons) or two-tailed student's t-test (two groups). The criterion for statistical significance was $P < 0.05$.

**References for Materials and Methods**

1. Z. Z. Xu et al., Inhibition of mechanical allodynia in neuropathic pain by TLR5-mediated A-fiber blockade. Nat.Med 21:1326-31 (2015).
2. T. Berta et al., Extracellular caspase-6 drives murine inflammatory pain via microglial TNF-alpha secretion. J Clin.Invest 124, 1173-1186 (2014).
3. G. Chen, C. K. Park, R. G. Xie, R. R. Ji, Intrathecal bone marrow stromal cells inhibit neuropathic pain via TGF-beta secretion. J Clin.Invest 125, 3226-3240 (2015).
4. H. Wakabayashi et al., Decreased sensory nerve excitation and bone pain associated with mouse Lewis lung cancer in TRPV1-deficient mice. J Bone Miner Metab 36, 274-285 (2018).
5. G. Chen et al., PD-L1 inhibits acute and chronic pain by suppressing nociceptive neuron activity via PD-1. Nature neuroscience 20, 917-926 (2017).
6. Q. Han et al., miRNA-711 Binds and Activates TRPA1 Extracellularly to Evoke Acute and Chronic Pruritus. Neuron 99, 449-463 e446 (2018).

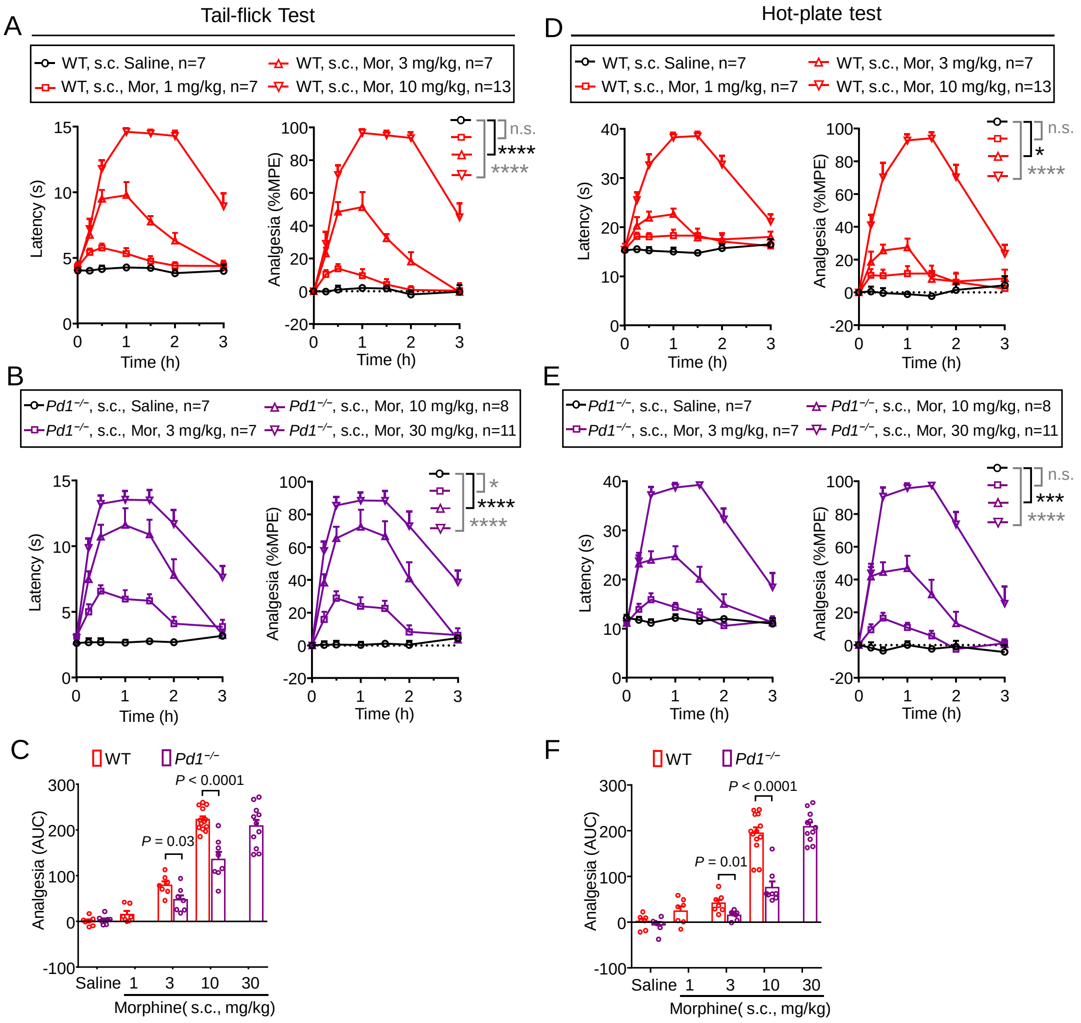


**Fig. S1. Dose-response of morphine analgesia in the WT mice and Pd1−/− mice following subcutaneous injection.** (**A-C**) Tail-flick test showing dose-response of morphine analgesia in WT and *Pd1*$^{-/-}$ mice. (A) Time course of tail-flick latency (left) and %MPE (right) of morphine antinociception (s.c., 1, 3, 10 mg/kg) in WT mice. *****P* < 0.0001, two-way ANOVA, followed by Dunnett's post hoc test, n = 7, 7, 7, 13 mice per group; n.s., not significant. (B) Time course of tail-flick latency (left) and %MPE (right) of s.c. morphine antinociception in *Pd1*$^{-/-}$ mic. **P* < 0.05, *****P* < 0.0001, two-way ANOVA, followed by Dunnett's post hoc test. n = 7, 7, 8, 11. (C) Comparison of AUC antinociception (%MPE) in WT mice (A) and KO mice (B). *P* = 0.03, *P* < 0.0001, two-tailed Student's t-test. (**D-F**) Hot plate test showing dose-response of morphine analgesia in WT and Pd1−/− mice. (D) Time course of hot-plate latency (left) and %MPE (right) of s.c. morphine in WT mice. **P* < 0.05. *****P* < 0.0001, two-way ANOVA, followed by Dunnett's post hoc test. n = 7, 7, 7, 13; n.s., not significant. (E) Time course of hot-plate latency (left) and %MPE (right) of s.c. morphine in *Pd1*$^{-/-}$ mice. ****P* < 0.001. *****P* < 0.0001, two-way ANOVA, followed by Dunnett's post hoc test. n = 7, 7, 8, 11 mice per group. (F) Comparison of AUC antinociception (%MPE) in WT mice (D) and KO mice (E). *P* = 0.01, *P* < 0.0001, two-tailed Student's t-test. Data are Mean ± SEM.

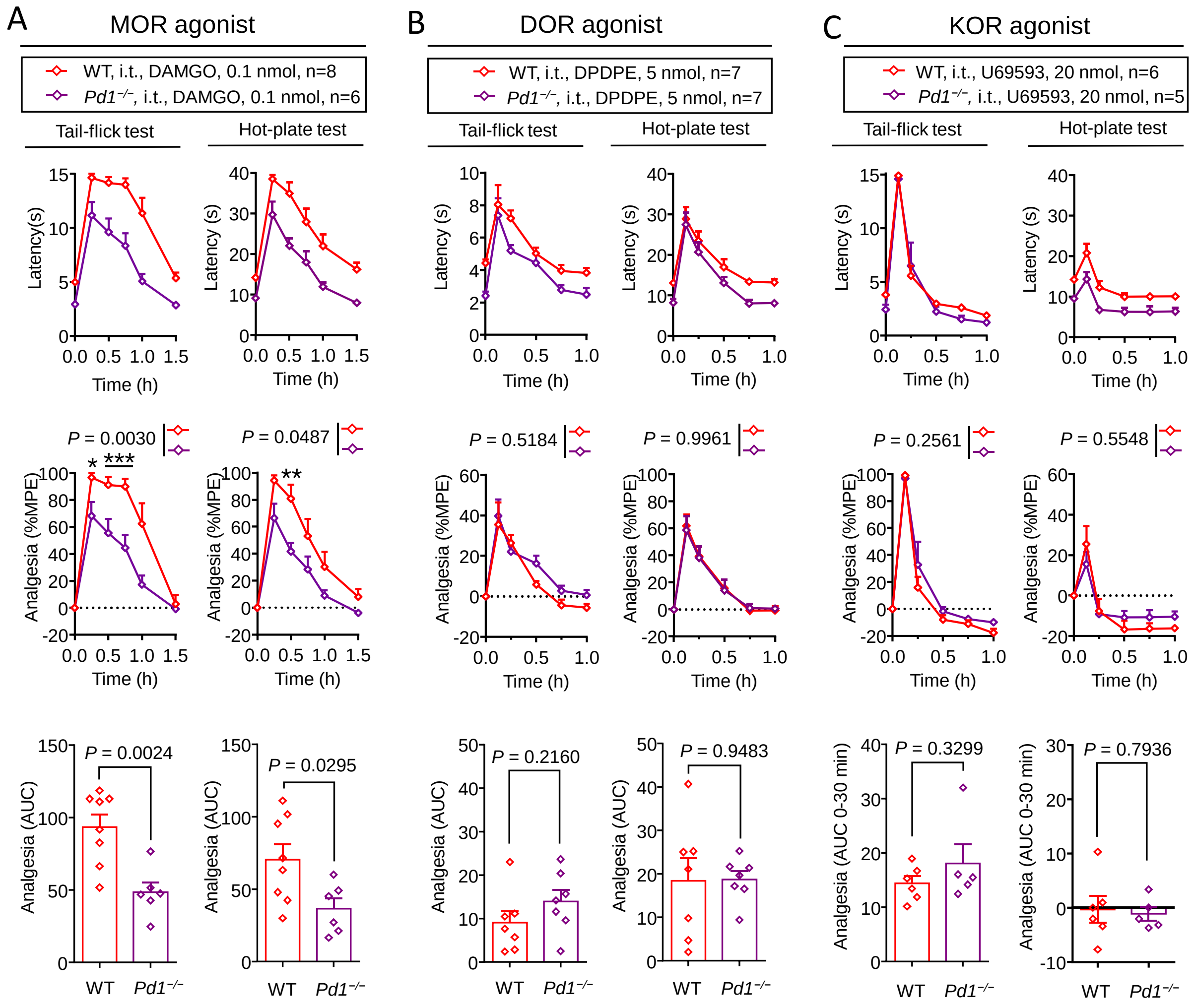


**Fig. S2**. **Antinociception of MOR selective agonist DAMGO, but not DOR agonist DPDPE or KOR agonist U69593 is decreased in *Pd1*$^{-/-}$ mice.** (**A**) Antinociception of MOR selective agonist DAMGO is decreased in *Pd1*$^{-/-}$ mice in both tail-flick test (left) and hot-plate test (right). Morphine antinociception was expressed as latency (top), %MPE (middle), and AUC of %MPE (bottom). For the %MPE data, $P$ = 0.0030, $P$ = 0.0487, two-way ANOVA; WT vs. KO, *$P$<0.05, **$P$<0.01, ***$P$<0.001, WT vs. KO, Bonferroni's post hoc test. For the AUC data, $P$ = 0.0024, $P$ = 0.0295, Two-tailed Student's t-test. n = 8 and 6 mice per group. (**B**) Antinociception of DOR selective agonist DPDPE is not altered in *Pd1*$^{-/-}$ mice in tail-flick test (left) and hot-plate test (right). For the %MPE data, $P$ = 0.5184, $P$ = 0.9961, WT vs. KO, two-way ANOVA, followed by Bonferroni's post hoc test. For the AUC data, $P$ = 0.2160, $P$ = 0.9483, WT vs. KO, two-tailed Student's t-test. n = 7 mice per group. (**C**) Antinociception of KOR selective agonist U69593 is not altered in *Pd1*$^{-/-}$ mice in both tail-flick test (left) and hot-plate test (right). For the %MPE data, $P$ = 0.5548, $P$ = 0.2561, WT vs. KO, two-way ANOVA, followed by Bonferroni's post hoc test. For the AUC data, $P$ = 0.3299, $P$ = 0.7935, WT vs. KO, Two-tailed Student's t-test. n = 6 and 5 mice per group. Note that U69593 produces no analgesia in the hot plate test in WT and KO mice. Data are Mean ± SEM.

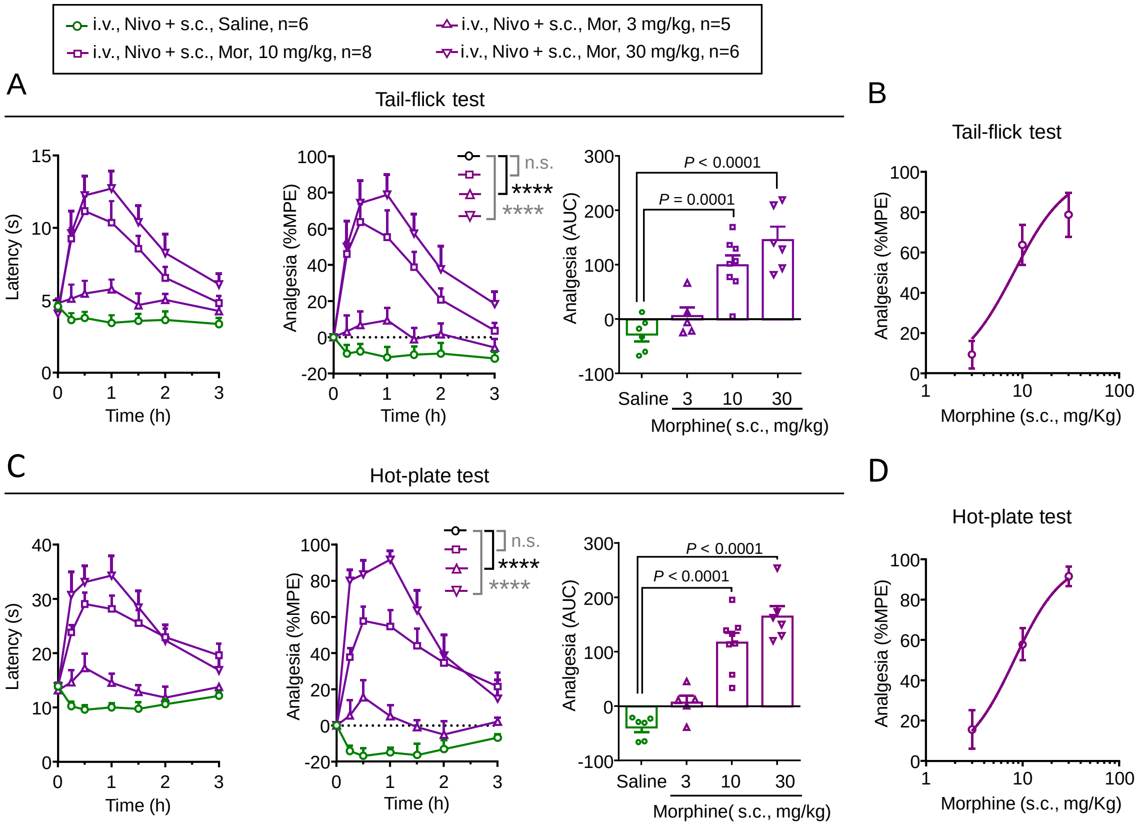


E

| ED $_{50}$ values (mg/kg) | WT | *Pd1*$^{-/-}$ | WT-Nivolumab pretreatment |
|---|---|---|---|
| Tail-flick test | 2.79<br>(2.32, 3.34) | 5.47<br>(3.80, 7.87) | 8.20<br>(5.29, 12.70) |
| Hot-plate test | 4.09<br>(3.44, 4.87) | 9.34<br>(8.06, 11.76) | 8.24<br>(6.08, 11.17) |

**Fig. S3. Dose-response of subcutaneous morphine analgesia in WT mice pretreated by intravenous Nivolumab (i.v., 10 mg/kg), given 30 min prior to the morphine injection.** (**A,B**) Tail-flick test showing dose-response of morphine analgesia (s.c., 1, 3, 10 mg/kg) in WT mice pretreated with Nivolumab. n = 6, 5, 8, 6 mice per group. (A) Time course of tail-flick latency. Left, tail-flick latency; Middle, %MPE; Right, AUC of %MPE. For the %MPE data, ****$P <$ 0.0001, vs. vehicle, two-way ANOVA; n.s., not significant. For the AUC data, $P$ = 0.0001, $P <$ 0.0001, one-way ANOVA, followed by Dunnett's post hoc test. (B) Dose-response curve of MPE at the peak time point. (**C, D**) Hot plate test showing dose-response of morphine analgesia (s.c., 1, 3, 10 mg/kg) in WT mice pretreated with Nivolumab. n = 6, 5, 8, 6 mice per group. (C) Time course of hot plate latency. Left, hot plate latency; Middle, %MPE; Right, AUC of %MPE. For the %MPE data, ****$P$<0.0001, vs. vehicle, two-way ANOVA; For the AUC data, $P <$ 0.0001, one-way ANOVA, followed by Dunnett's post hoc test. (D) Dose-response curve of MPE at the peak time point. (**E**) The ED50 values of the antinociceptive effects of s.c. injection of morphine in the WT mice and *Pd1*$^{-/-}$ mice (Fig. 1E) and Nivolumab pretreated mice (B, D). Data are Mean ± SEM.

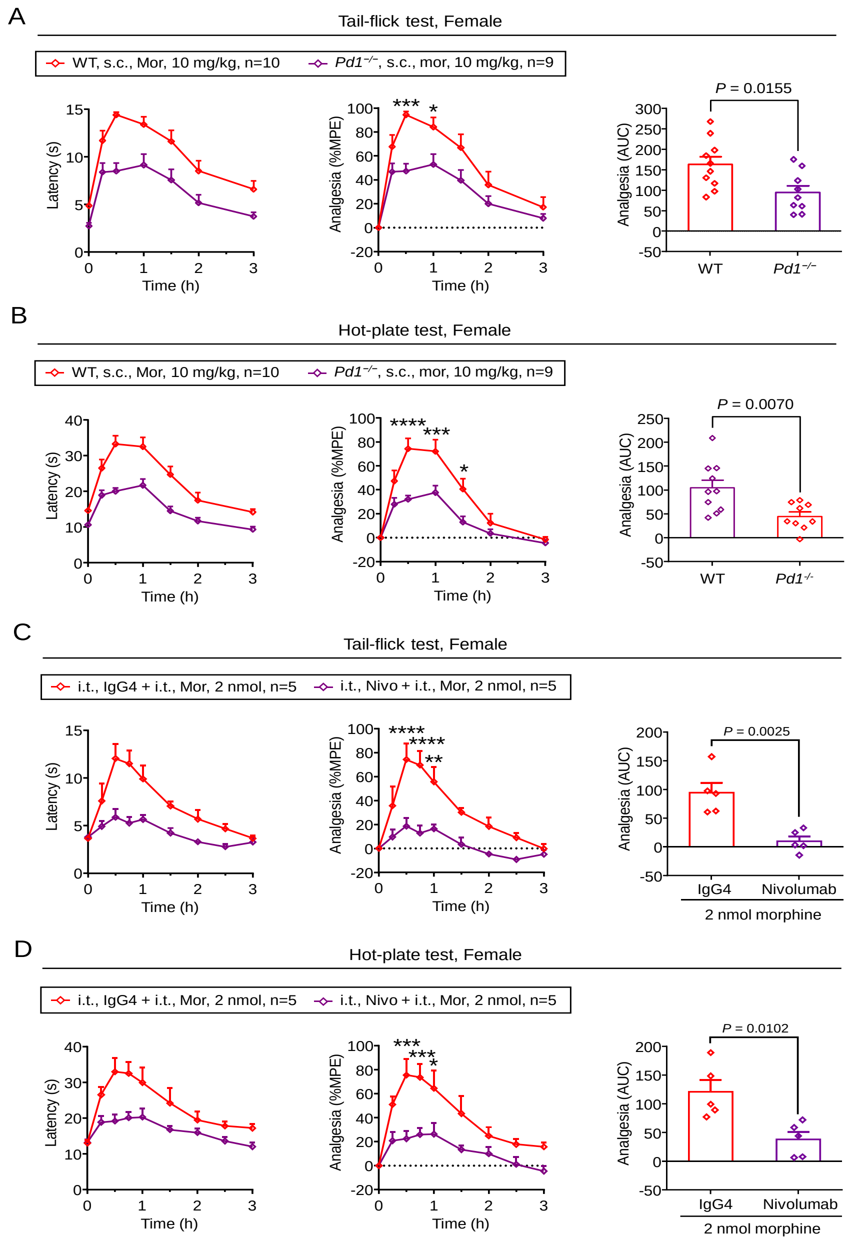


**Fig. S4. *Pd1* deficiency and Nivolumab pretreatment inhibit morphine antinociception in female mice.** (**A, B**) Tail-flick test (A) and hot plate test (B) showing morphine analgesia (s.c., 10 mg/kg) in WT and *Pd1*$^{-/-}$ female mice. Left, Tail-flick latency; Middle, %MPE; Right, AUC of %MPE, *P* = 0.0155 (A), *P* = 0.0070 (B), two-tailed Student's t-test. WT vs. KO, n = 10 and 9 mice per group. (**C, D**) Tail-flick test (C) and hot plate test (D) showing morphine analgesia (i.t., 2 nmol) in WT mice pretreated with Nivolumab (Nivo) or control IgG4 (i.t., 1 µg), given 30 min prior to the morphine injection. Left, Tail-flick latency; Middle, %MPE; Right, AUC of %MPE, *P* = 0.0025 (c), *P* = 0.0102 (d), two-tailed Student's t-test. WT vs. KO, n = 5 mice per group. Data are Mean ± SEM. **P* < 0.05, ***P* < 0.01, ****P* < 0.001, *****P* < 0.0001, vs. KO or vs. Nivo, two-way ANOVA, followed by Bonferroni's post hoc test.

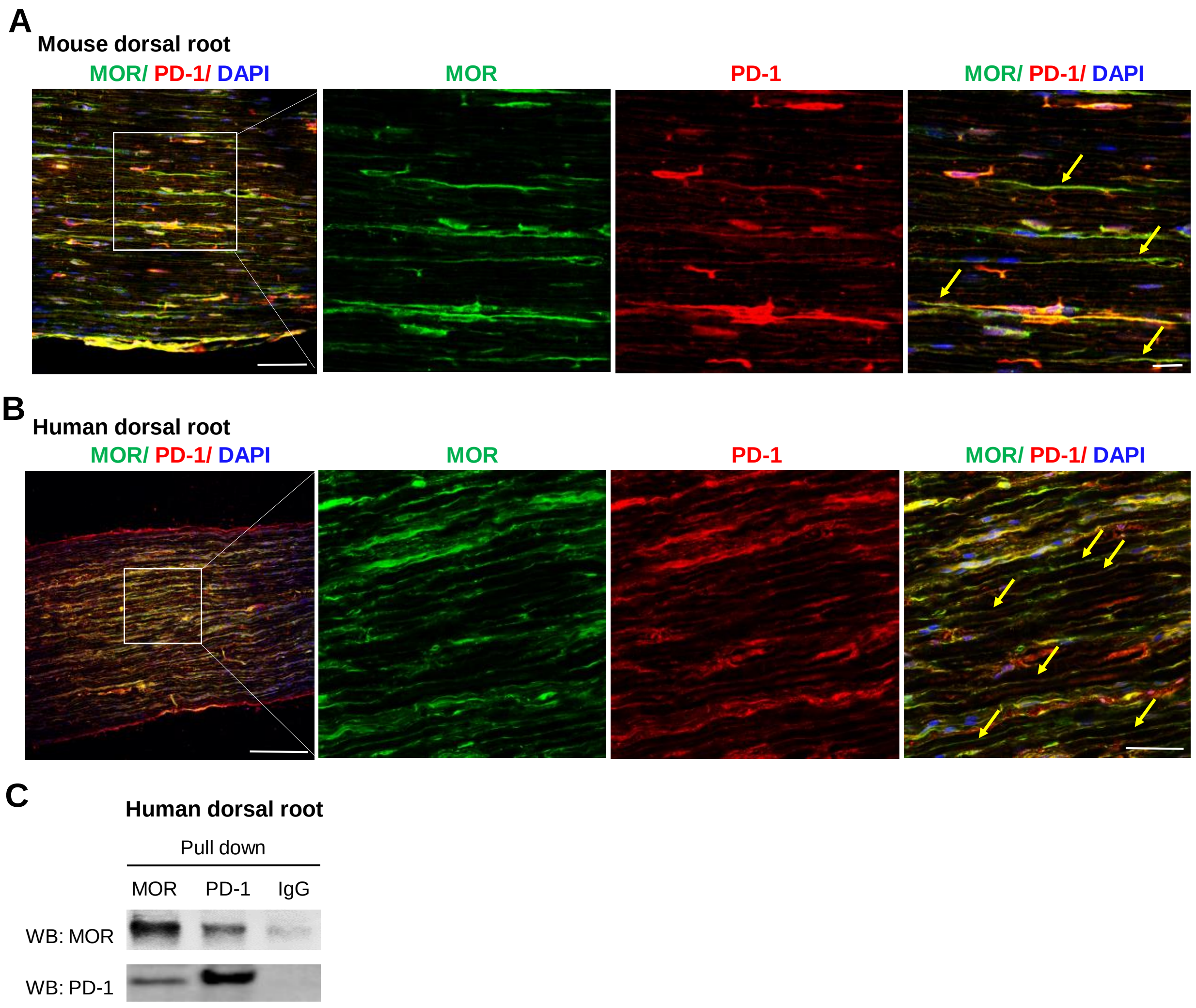


**Fig. S5. PD-1 and MOR are co-expressed in axons of mouse and human dorsal roots.** (**A**) Double staining for MOR and PD-1 in mouse dorsal root. Arrows indicate double-labeled nerve fibers. Scale bars, 75 μm (left) and 20 μm (right). (**B**) Double staining for MOR and PD-1 in human dorsal root. PD-1 was labeled by Nivolumab (1 mg/ml). Arrows indicate the double-labeled nerve fibers. Scale bars, 250 μm (left) and 50 μm (right). (**C**) Co-IP showing PD-1/MOR interaction in human dorsal root tissue. Human dorsal root lysates were immunoprecipitated with MOR or PD-1 antibody and then immunoblotted with PD1 or MOR antibody.

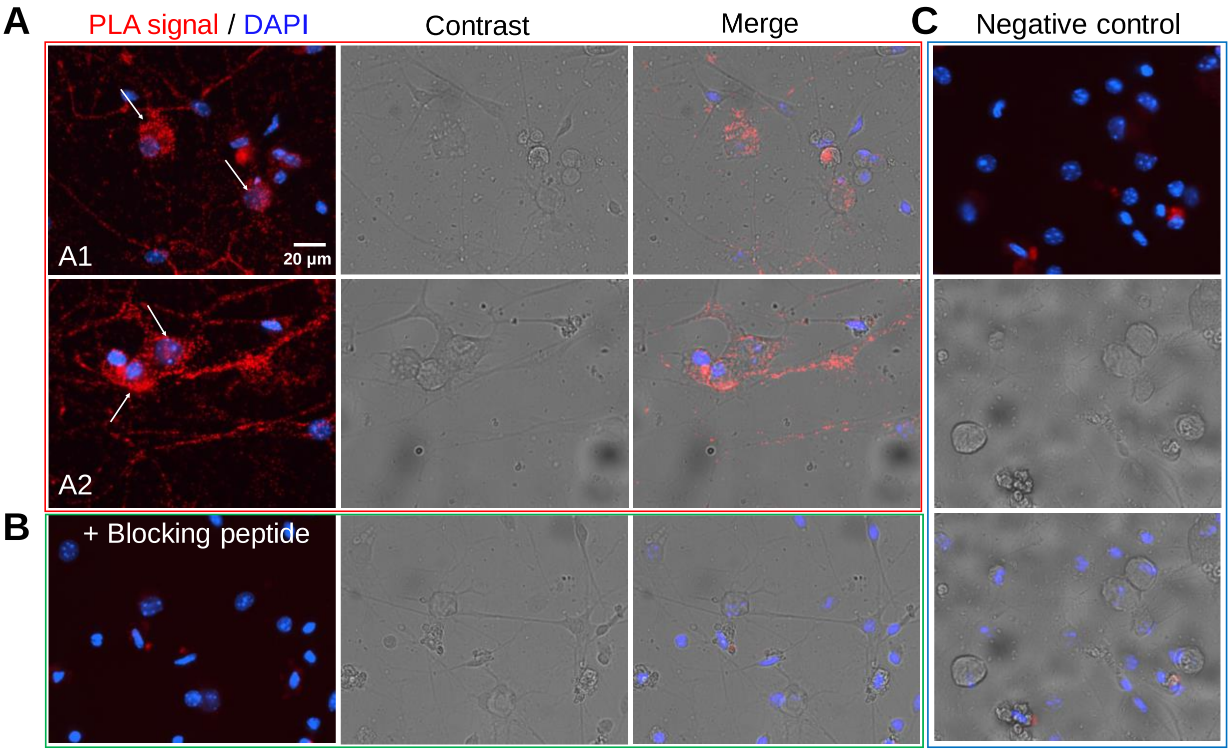


**Fig. S6. Proximity ligation assay (PLA) reveals PD-1/MOR interaction in cultured mouse DRG neurons. (A)** Positive PLA labeling in two DRG cultures (A1 and A2). Contrast images show all the cells in two optic fields. Arrows indicate positive neurons. Scale bar, 20 µm. (**B**) PLA signal is blocked by adding the blocking peptide for the PD-1 antibody (1:200, Sigma, catalog: SBP4065). (**C**) Negative control in the absence of primary antibodies. PLA was performed using Duolink reagents (Sigma-Aldrich, catalog: DUO92101) on cultured mouse DRG neurons. Cells were incubated with a mixture of anti-PD1 antibody (1:100 rabbit) and anti-MOR antibody (1:200 mouse) overnight at 4°C. Then the cells were incubated with respective secondary antibodies (anti-mouse MINUS probe and anti-rabbit PLUS probe), followed by incubation with the polymerase in amplification buffer for 100 min at 37 °C.

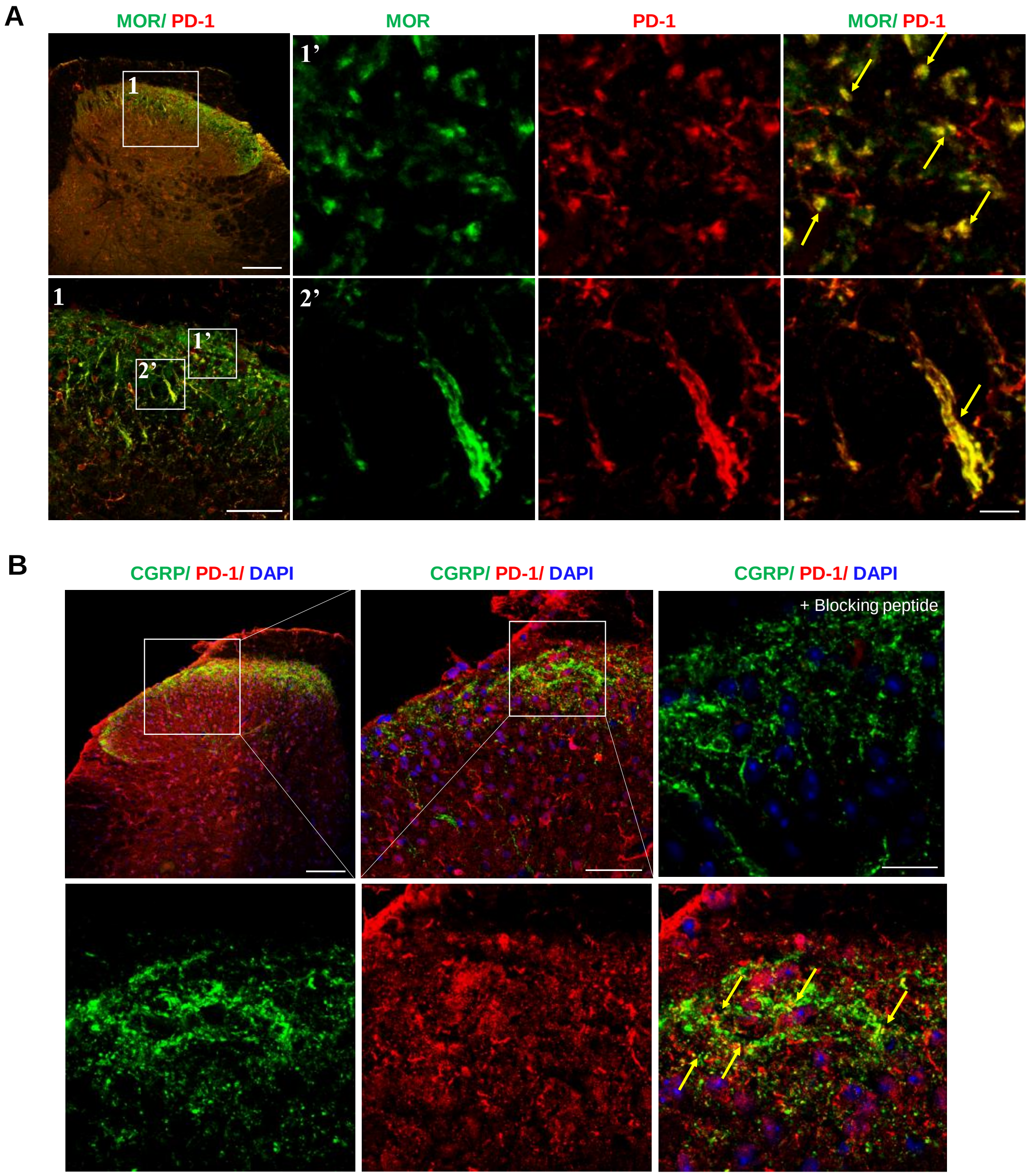


**Fig. S7. PD-1 is co-expressed with MOR and CGRP in axonal terminals in mouse spinal dorsal horn.** (**A**) Double staining of PD-1 (red) and MOR (green) in axonal terminals in the spinal dorsal horn. Top left, the small box is enlarged in the bottom left panel, in which two boxes (1’ and 2’) are further enlarged in the right panels (top for 1’ and bottom for 2”). Three separate images in 1’ and 2’ are single labeling and merged images. Scale bars, 100 µm (top left), 50 µm (bottom left), and 20 µm (right). (**B**) Double staining for PD-1 (red) and CGRP (green) in axonal terminals in spinal dorsal horn. Top left, the small box is enlarged in the right panels. Top middle, the small box is enlarged in three separate boxes with single labeling and merged images in the bottom panels. Scale bars, 100 µm (left), 50 µm (medium), and 20 µm (right). Arrows indicate the double-labeled axonal terminals.

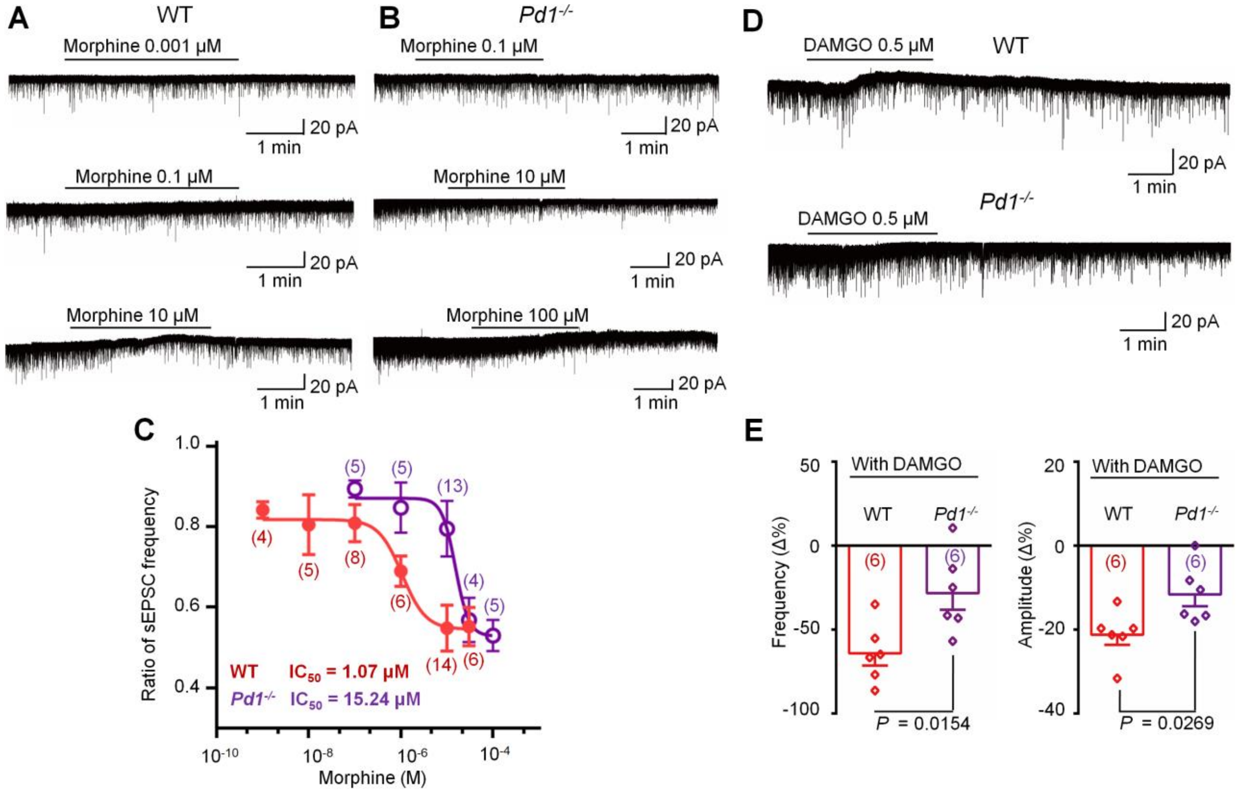


**Fig. S8. Effects of morphine and DAMGO on sEPSCs in lamina IIo neurons of spinal cord slice in WT and *Pd1*$^{-/-}$ mice.** (**A,B**) Traces of sEPSCs before and after morphine treatment in WT and *Pd1*$^{-/-}$ mice at different concentrations. (**C**) Dose response curves of morphine inhibition of sEPSCs in WT and KO mice. Note the $IC_{50}$ for morphine inhibition of sEPSC is 1.07 μM and 15.24 μM in WT and KO mice, respectively. The dose-response curves were drawn according to the Hill equation. Number of neurons in each group is indicated in respective bracket. (**D**) Traces of sEPSCs before and after DAMGO treatment in WT and *Pd1*$^{-/-}$ mice. (**E**) DAMGO-induced reduction in sEPSC frequency (left) and amplitude (right) in WT and KO mice. n=6 neurons per group, unpaired two-tailed t-test. Number of neurons in each group is indicated in respective bracket. Data are Mean ± SEM.

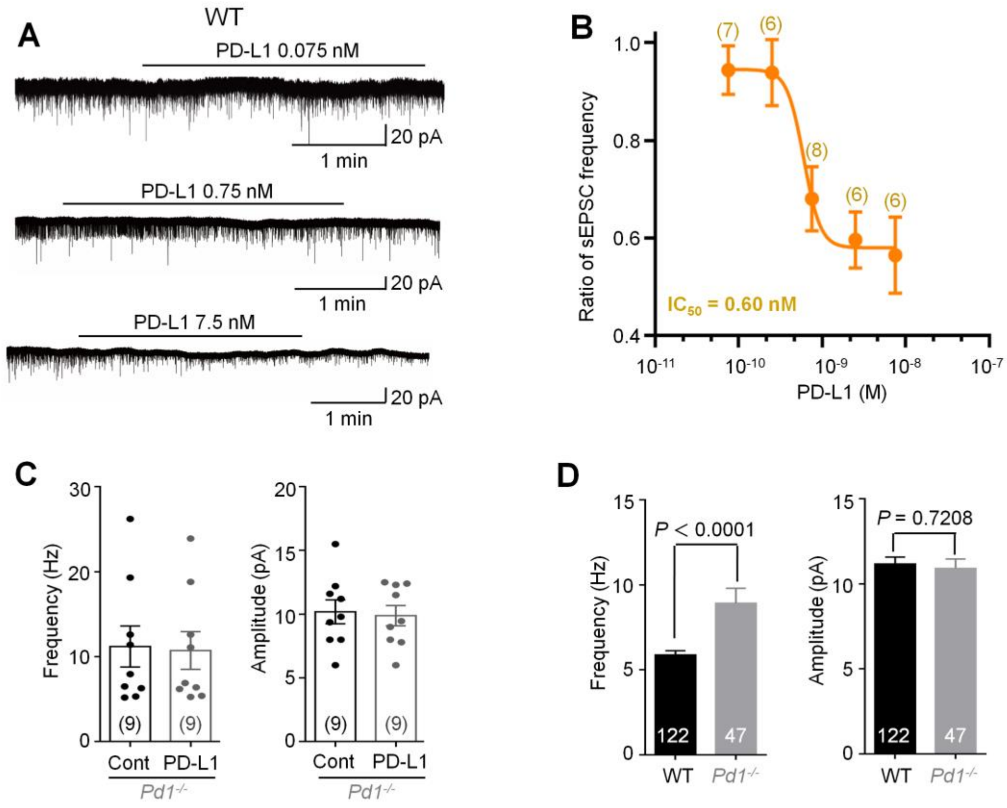


**Fig. S9. Effects of PD-L1 on sEPSCs in lamina IIo neurons of spinal cord slice in WT and *Pd1*$^{-/-}$ mice.** (**A**) Traces of sEPSCs before and after PD-L1 treatment in WT mice. (**B**) Dose response curves of PD-L1 inhibition of sEPSC frequency in WT mice. Note the $IC_{50}$ for PD-L1's inhibition of sEPSC (0.60 nM) is much lower than that of morphine (1.07 μM). (**C**) sEPSC frequency (left) and amplitude (right) before (control, Cont) and after PD-L1 treatment (7.5 nM) in WT and KO mice, n=9 cells per group. (**D**) sEPSC frequency (left) and amplitude (right) in WT (n=122 cells) and KO mice (n=47 cells), unpaired two-tailed t-test. Data are Mean ± SEM.

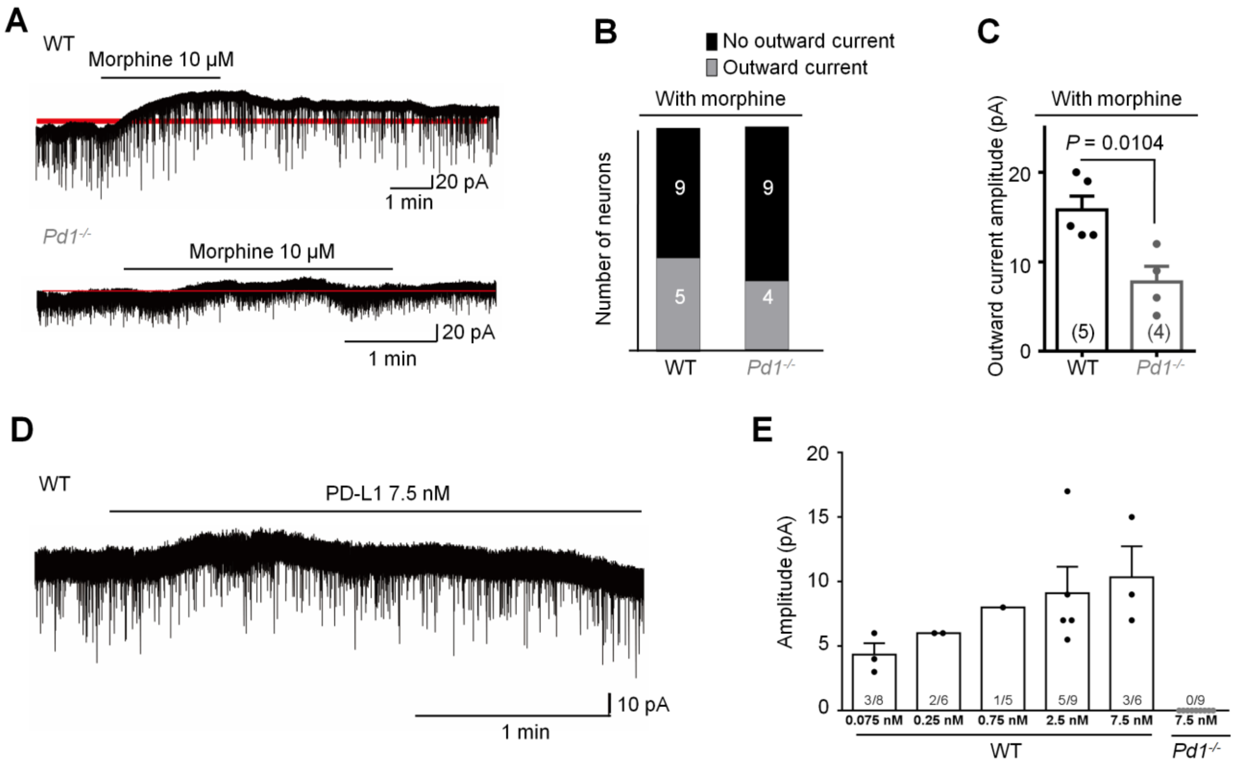


**Fig. S10. Morphine and PD-L1 induce outward currents in lamina II neurons of spinal cord slices in WT but not *Pd1*$^{-/-}$ mice.** (**A,B**) Traces of Morphine-induced outward currents in WT and *Pd1*$^{-/-}$ mice. (**B**) Number of neurons with and without outward currents in WT and *Pd1*$^{-/-}$ mice. (**C**) Amplitude of morphine-induced outward currents in WT and *Pd1*$^{-/-}$ mice (n = 5 in WT and n = 4 in *Pd1*$^{-/-}$ mice; *P* = 0.0104, unpaired two-tailed t-test). (**D**) Trace of PD-L1 (7.5 nM) induced outward current. (**E**) Amplitude of PD-L1-induced outward currents in WT and *Pd1*$^{-/-}$ mice. Number of total neurons and responding neurons is indicated in each column. Data are Mean ± SEM.

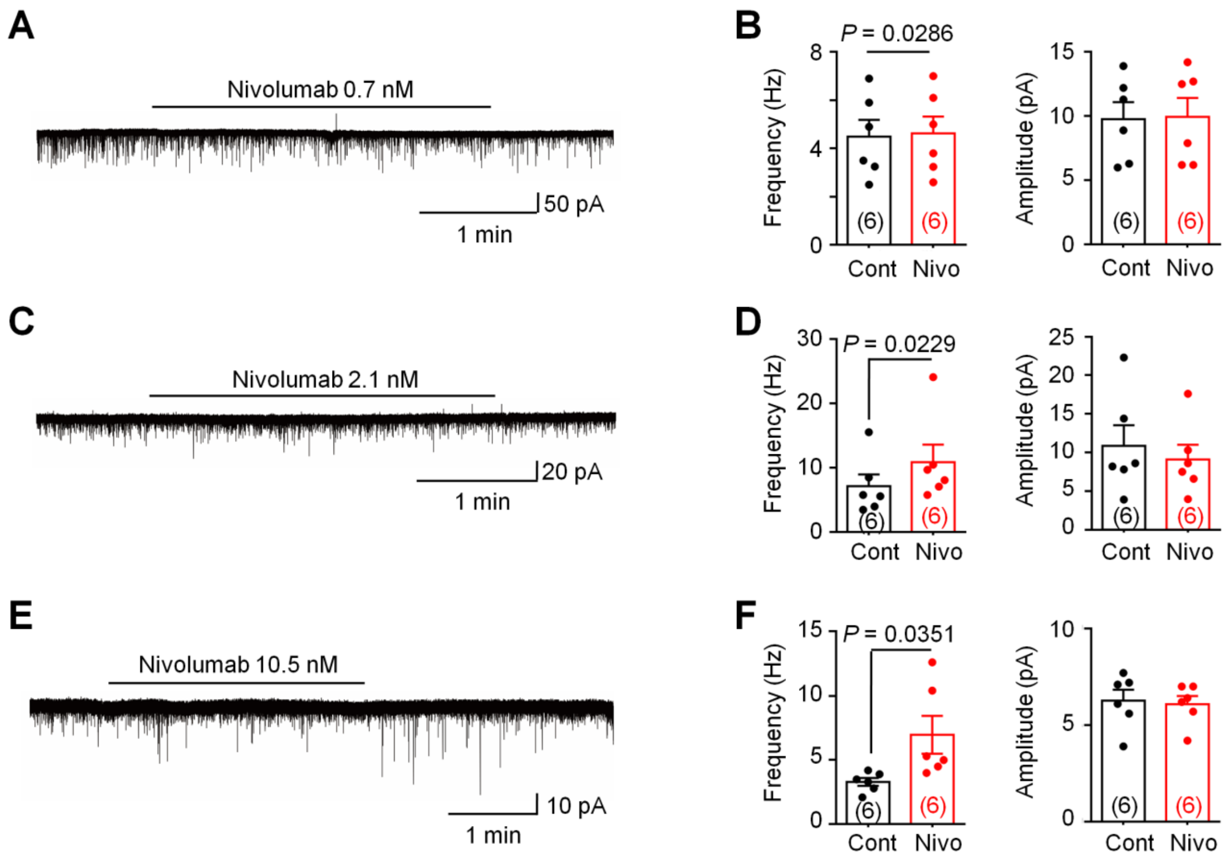


**Fig. S11. Anti-PD-1 treatment with Nivolumab increases sEPSC frequency in lamina II neurons of spinal cord slices.** (**A,C,E**) Traces of sEPSCs before and after Nivolumab treatment at different concentrations. (**B,D,F**) Frequency (left) and amplitudes (right) of sEPSCs before and after Nivolumab treatment, unpaired two-tailed t-test. n=6 neurons per group. The same concentration of Nivolumab was used in A and B, C and D, and E and F. Data are Mean ± SEM.

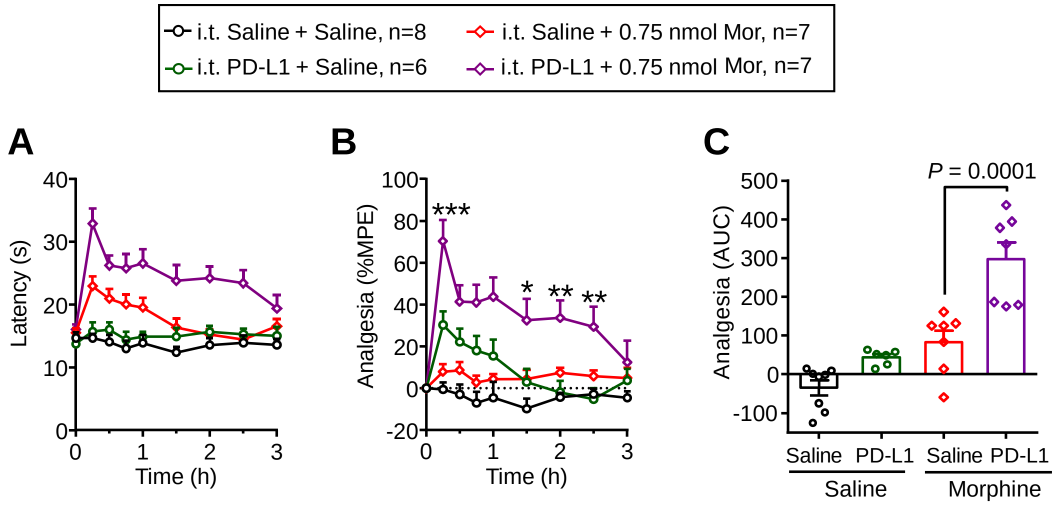


**Fig. S12. Spinal antinociception of morphine following i.t. injection is enhanced by i.t. pretreatment with PD-L1 in hot-plate test.** (**A-C**) Hot-plate test showing enhanced morphine analgesia by PD-L1. Spinal antinociception of morphine following i.t. injection is enhanced by i.t. pretreatment with PD-L1 in the. PD-L1 (3 µg) or saline was i.t. injected 30 min prior to morphine injection (i.t., 0.75 nmol). (A) Time course of hot plate latency. (B) Time course of spinal antinociception (%MPE), **P* < 0.05, ***P* < 0.01, ****P* < 0.0001, vs. saline pretreatment control, two-way ANOVA, followed by Bonferroni's post hoc test. (C) AUC of %MPE from b. P = 0.0001, vs. saline pretreatment group, one-way ANOVA, followed by Bonferroni's post hoc test. n = 8, 6, 7, 7 mice per group. Data are Mean ± SEM.
